# Early cis-regulatory events in the formation of retinal horizontal cells

**DOI:** 10.1101/2020.12.02.409003

**Authors:** Estie Schick, Kevin C. Gonzalez, Pooja Dutta, Kazi Hossain, Miruna G. Ghinia Tegla, Mark M. Emerson

## Abstract

During retinal development, multipotent and restricted progenitor cells generate all of the neuronal cells of the retina. Among these are horizontal cells, which are interneurons that modulate the light-induced signal from photoreceptors. This study utilizes the identification of novel cis-regulatory elements as a method to examine the gene regulatory networks that direct the development of horizontal cells. Here we describe a screen for cis-regulatory elements, or enhancers, for the horizontal cell-associated genes PTF1A, ONECUT1 (OC1), TFAP2A (AP2A), and LHX1. The OC1ECR22 and Tfap2aACR5 elements were shown to be potential enhancers for OC1 and TFAP2A, respectively, and to be specifically active in developing horizontal cells. The OC1ECR22 element is activated by PTF1A and RBPJ, which translates to regulation of OC1 expression and suggests that PTF1A is a direct activator of OC1 expression in developing horizontal cells. The region within the Tfap2aACR5 element that is responsible for its activation was determined to be a 100 bp sequence named Motif 4. Both OC1ECR22 and Tfap2aACR5 are negatively regulated by the nuclear receptors THRB and RXRG, as is the expression of OC1 and AP2A, suggesting that nuclear receptors may have a role in the negative regulation of horizontal cell development.

## Introduction

The vertebrate retina is comprised of six major classes of neuronal cells. Cone and rod photoreceptors (PRs) receive a light stimulus and, through the phototransduction cascade, convert it into an electrical signal which is passed on to bipolar cells (BCs). The BCs ultimately send the impulse to retinal ganglion cells (RGCs), whose axons form the optic nerve. Horizontal cells (HCs) and amacrine cells (ACs) are interneurons that participate in the transmission and modulation of this signal. HCs form lateral connections to provide inhibitory feedback to photoreceptors, while ACs regulate bipolar cell output to RGCs. These cells are born in an overlapping chronological order that is conserved among many vertebrates (Bassett and Wallace, 2012). While the proportions of these cell types are varied across species, the cells are organized in a conserved retinal structure with three nuclear layers separated by two plexiform layers where synaptic connections are made.

HCs are a highly-specialized class of inhibitory interneurons that possess several unique features that distinguish them from other retinal cell types. Firstly, HCs have been shown to migrate past their prospective layer in the retina before settling in their final position in the outer portion of the inner nuclear layer (INL) (Edqvist and Hallböök, 2004) In addition, there have been several reports of dedicated precursors that divide symmetrically to give rise to two HCs (Godinho et al., 2007; Rompani and Cepko, 2008). In zebrafish, most HC precursors migrate toward their position in the INL before the final mitosis (Amini et al., 2019; Godinho et al., 2007), while in chick, the final mitosis of HC precursors takes place at the basal membrane before subsequent migration toward the INL (Boije et al., 2009). Furthermore, several HC subtypes are present in the retina and can be readily distinguished by morphology and expression of molecular markers (Fischer et al., 2007). All vertebrates possess HCs (Boije et al., 2016), although the number of subtypes varies across species. Avian species are reported to have 2-4 HC types (Fischer et al., 2007; Mariani and Leure-DuPree, 1977), while only 1 HC subtype has been identified in the mouse retina (Peichl and González-Soriano, 1994) and 3 HC types have been reported in the human retina (Ahnelt and Kolb, 1994; Kolb et al., 1992, 1994).

The gene regulatory network (GRN) that underlies and directs the development of HCs has yet to be fully elucidated. FOXN4, a winged helix/forkhead transcription factor (TF), has been shown to be a key factor for the fate commitment of HCs, and accordingly, there is a complete absence of HCs in the FOXN4 knockout (KO) (Li et al., 2004). A downstream target of FoxN4 is the bHLH factor PTF1A, whose loss leads to the same HC phenotype (Fujitani et al., 2006). These factors are both necessary and sufficient for the generation of HCs but are similarly required for the formation of ACs, which are drastically decreased in the FOXN4 and PTF1A knockouts (Fujitani et al., 2006; Li et al., 2004). An additional series of TFs are therefore necessary to specify the HC fate. Several other genes have been reported to be important for HC development, which include ONECUT1 (OC1), TFAP2A/B (AP2A), and LHX1 (LIM1). More than 80% of HCs fail to develop upon the conditional KO of OC1 (Wu et al., 2013) and the double KO of OC1 and OC2 results in the complete loss of HCs (Sapkota et al., 2014). TFAP2A is reported to be a downstream effector of PTF1A, and the double KO of TFAP2A and TFAP2B results in a near complete loss of HCs (Bassett et al., 2012; Jin et al., 2015). Finally, LHX1 has been shown to be essential for the proper lamination of HCs, as these cells are ectopically located in the LHX1 KO (Poché et al., 2007). These transcription factors, among others, therefore play a role in the GRN that directs the development of HCs.

While there are several proposed models for HC development, many relationships between genes are drawn upon downregulation in a knockout environment, without evidence of direct activation. Knowledge of the cis-regulatory modules (CRMs), or enhancer elements, that regulate gene expression can add a level of resolution to current models of development. These elements will provide a foothold for the study of the GRN that directs HC development, as well as serve as a much-needed tool to target developing HCs. In this study, we sought to detect CRMs for PTF1A, ONECUT1, TFAP2A and LHX1 that are specifically active in HCs and to determine the transcription factors that occupy them. In doing so, we identified OC1ECR22 and Tfap2aACR5 as novel enhancer elements that display preferential activity in HCs in addition to other elements that, while not immediately relevant to this study, are active in specific cell populations. We detected conserved binding sites for PTF1A and RBPJ within the OC1ECR22 sequence and observed that mutation of these sites resulted in the loss of all activity of the enhancer, implicating PTF1A and RBPJ in direct activation of the OC1ECR22 element and OC1 expression. We identified a 100 bp region necessary for the activation of the Tfap2aACR5 element, and finally, we uncovered a potential role for nuclear receptors in negatively regulating HC development.

## Results

### Screen for enhancer elements near genes involved in horizontal cell development

To uncover the mechanisms that underlie the generation of HCs, we sought to identify CRMs with activity in HCs. We developed a screen for regulatory elements in proximity to the PTF1A, TFAP2A, LHX1, and OC1 loci, as these genes are known to be involved in HC development. We identified both evolutionary conserved regions (ECRs) with ECR Browser and accessible chromatin regions (ACRs) with an ATAC-Seq dataset previously reported by our lab (Patoori et al., 2020) (Figure 1A). Although elements were named as ECRs or ACRs based on the method of their identification, some ECRs are located within accessible chromatin and some ACRS are conserved evolutionarily. With these analyses, we identified 40 candidate enhancer elements and isolated them from the chicken genome (Table 1; this includes three OC1-associated elements originally identified in Patoori et al., 2020). The DNA elements were cloned into an expression plasmid upstream of a minimal promoter element and GFP and PLAP reporter genes, enabling us to test their ability to drive reporter activity in the developing chick retina (Figure 1B). Retinas from embryonic day 5 (E5) chick embryos were electroporated with one of these plasmids and a broadly active CAG::mCherry plasmid as an electroporation control. Expression of the AP reporter after two days in culture was used as a reporter of enhancer activity. Varying levels of enhancer-driven activity were present with 19 of the DNA elements; this was observed even among elements near the same gene. For example, among 4 elements adjacent to PTF1A, both the level and location of activity were varied (Figure 1C, Figure S1). The activity of these putative enhancers was confirmed with the GFP reporter. Furthermore, we counterstained retinas for Visinin, a marker of PRs, in order to eliminate candidate enhancers with substantial activity in that population. For example, the activity driven by Tfap2aACR2 does not overlap with Visinin expression and is therefore a promising candidate, whereas activity driven by Lhx1ACR2 and Ptf1aACR1 was present in cells that express Visinin and were eliminated from the screen (Figure 1D). In total, enhancer-driven activity with 10 of the elements was observed primarily in the inner retina, where migrating HCs are localized at this time point (Boije et al., 2009) (Figure 1D, Figure S2).

**Figure 1.**
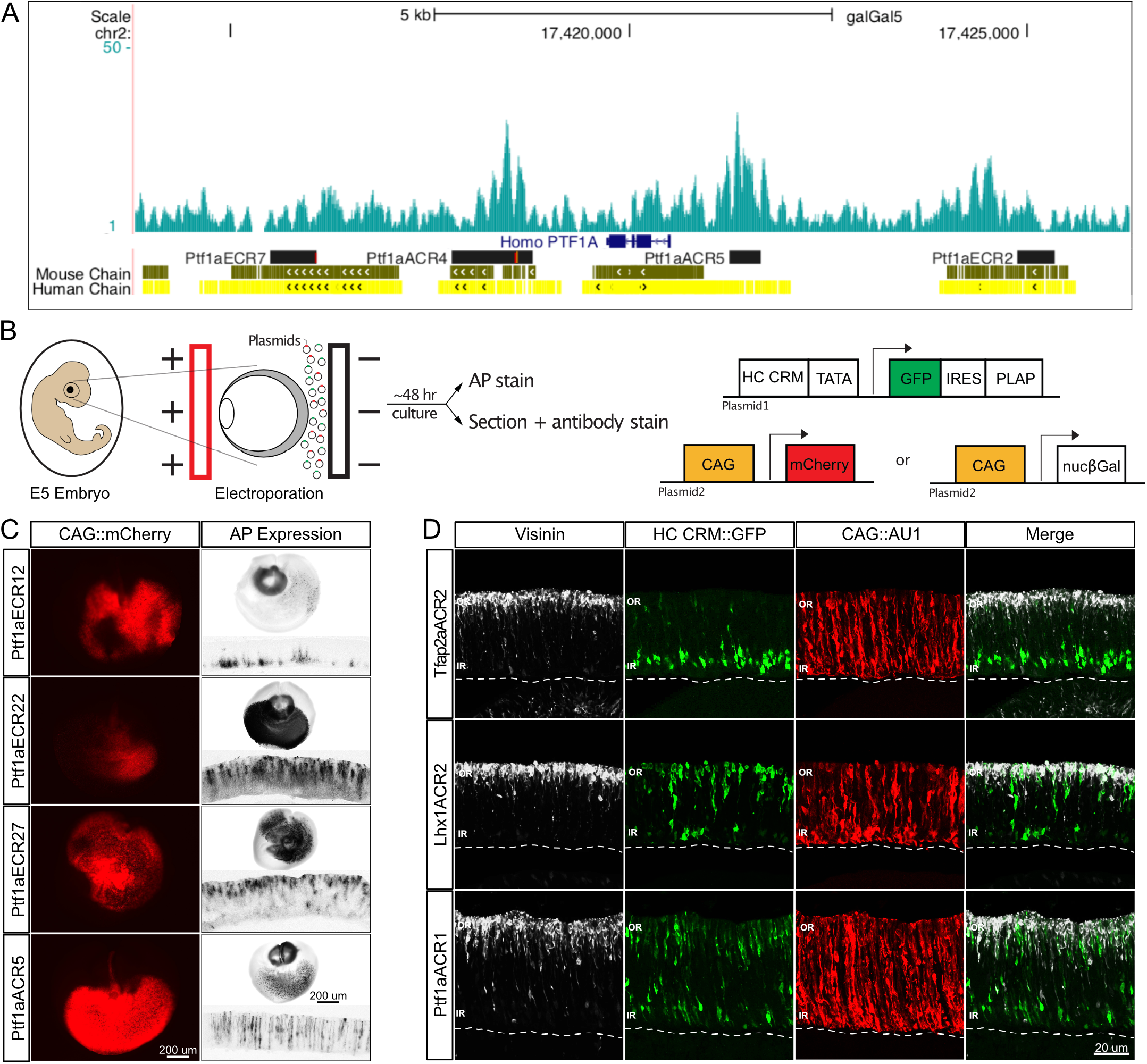
Active cis-regulatory elements near horizontal cell-associated genes have varied levels and patterns of activity. A. Schematic of experimental workflow. B. View of ATAC-Seq reads aligned to the galGal5 genome in UCSC Genome Browser at the PTF1A locus. Peaks indicate chromatin accessibility, black rectangles represent selected DNA elements. Conservation of chick sequences to mouse and human genomes represented by Mouse Chain and Human Chain tracks. C. AP reporter assay for HC CRM activity. Representative images of whole retinas electroporated *ex vivo* at E5 with CAG::mCherry as an electroporation control and Ptf1a CRM::PLAP. After 2 days in culture, AP stain was developed. Scale bars represent 200 μm. D. Visinin counterstain rules out HC CRMs with activity in photoreceptors. Representative images of vertically sectioned retinas that were electroporated *ex vivo* at E5 with CAG::AU1 as an electroporation control and HC CRM::GFP. After 2 days in culture, retinas were stained for Visinin. Maximum intensity projection of 40x image, scale bars represent 20 μm. ECR, evolutionary conserved region; ACR, accessible chromatin region; AP, alkaline phosphatase; CRM, cis-regulatory module; OR, outer retina; IR, inner retina.

Because enhancers are active transiently during development, this AP and GFP expression is only representative of cells that experienced enhancer activity near the time of analysis. In order to obtain a more accurate assessment of the cells with enhancer activity, we utilized a lineage tracing system to convert reporter activity into a constitutive signal upon initial activity of the DNA element and thereby provide a history of enhancer activity. This system has previously been shown to have no leaky expression in control conditions (Schick et al., 2019). The 10 elements with activity in a HC-like pattern were cloned into lineage tracing plasmids and electroporated into E5 chick retinas alongside CAG::nβgal as an electroporation control (Figure 2A). After 2 days in culture, we counterstained the retinas for Visinin or for LIM1, a specific marker of some HCs. The lineage tracing revealed three patterns of enhancer-driven activity: 1) some elements, such as Lhx1ACR1, drove reporter expression non-specifically throughout the retina; 2) others, such as Tfap2aACR4, drove similar amounts of reporter expression in the outer and inner limits of the retina; 3) the remaining elements, such as Tfap2aACR5, drove reporter expression largely in the inner retina (Figure 2B-C). This last category is comprised of Ptf1aECR12, Tfap2aACR2, Tfap2aACR5, OC1ECR22, and OC1ACR8. These 5 elements drove reporter expression in cells organized in the inner retina that did not have Visinin expression, and that showed some colocalization with LIM1 (Figure 2B-C, Figure S3). This suggests that these elements are specifically activated in cells that will become HCs and not photoreceptors.

**Figure 2.**
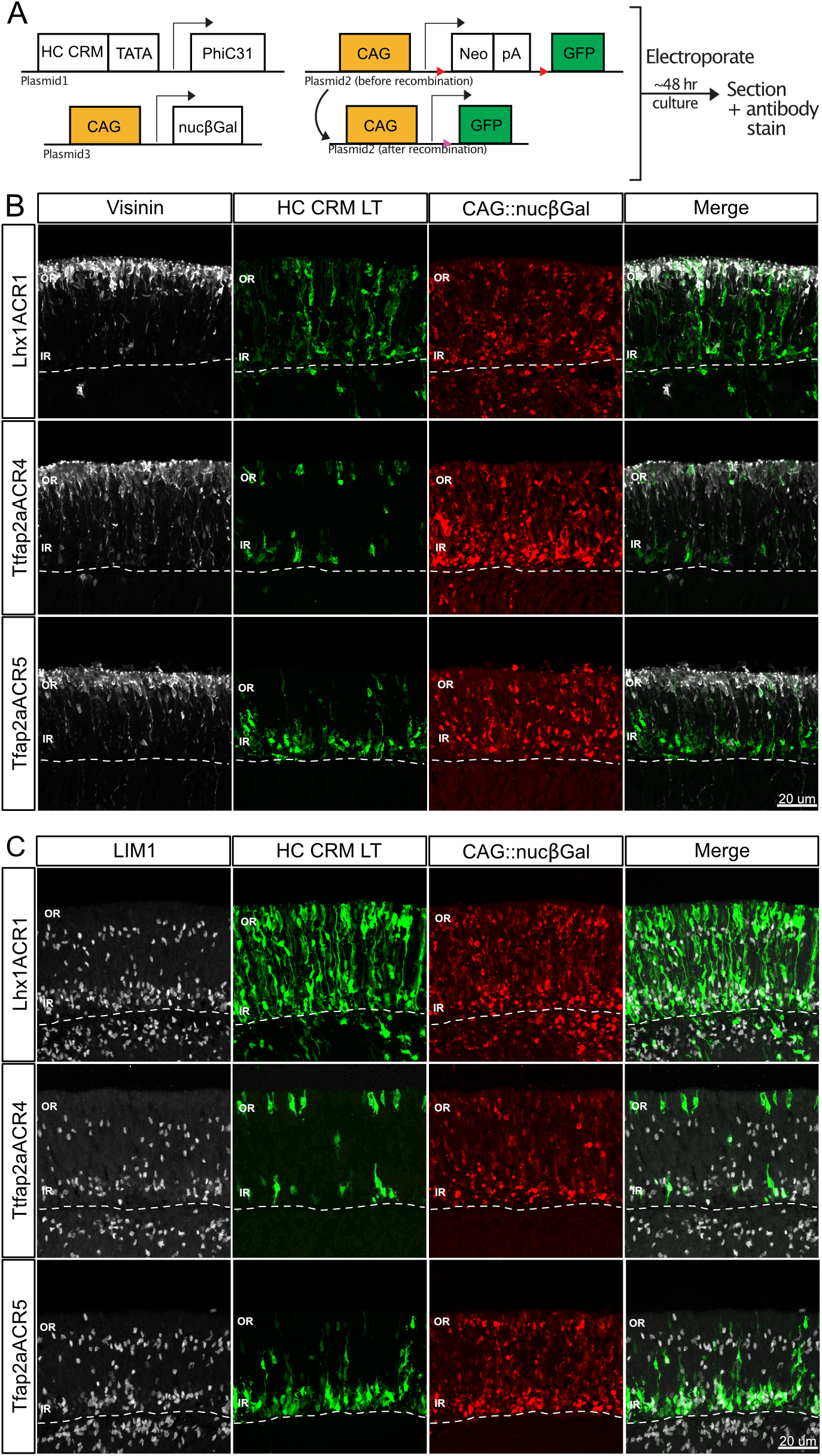
Lineage tracing of active CRMs allows more extensive characterization. A. Schematic of experimental workflow. B-C. Visinin and LIM1 highlight HC CRM activity in photoreceptors and horizontal cells, respectively. Representative images of vertically sectioned retinas that were electroporated *ex vivo* at E5 with CAG::nucβgal as an electroporation control, HC CRM::PhiC31 and CAaNa::GFP. After two days in culture, retinas were counterstained for Visinin (B) or LIM1 (C). Maximum intensity projection of 40x image, scale bars represent 20 μm. CRM, cis-regulatory module; pA, poly adenylation; OR, outer retina; IR, inner retina.

### Characterization of enhancer-driven activity

We further characterized the activity of these elements by electroporating the lineage tracing plasmids into E5 chick retinas and dissociating the cells after three days in culture. We then counterstained the cells for a variety of markers for all cell types in the retina, and using flow cytometry, we quantified the percentage of cells with expression of each marker among the population labeled with GFP (Figure 3A). We first stained the dissociated cells for LIM1 and AP2A. LIM1 is expressed in H1 HCs, which represent 52% of HCs in the chick, and AP2A marks those HCs as well as ACs (Fischer et al., 2007). Therefore, the cells with expression of both LIM1 and AP2A were identified as HCs, while the cells with expression of AP2A alone were considered to be ACs. All 5 enhancers are active in H1 HCs, which represent a range of 14-26% of cells with enhancer activity (Figure 3B). Ptf1aECR12 had substantial activity in ACs, while ACs represent only 1-7% of the other enhancer populations (Figure 3C).

**Figure 3.**
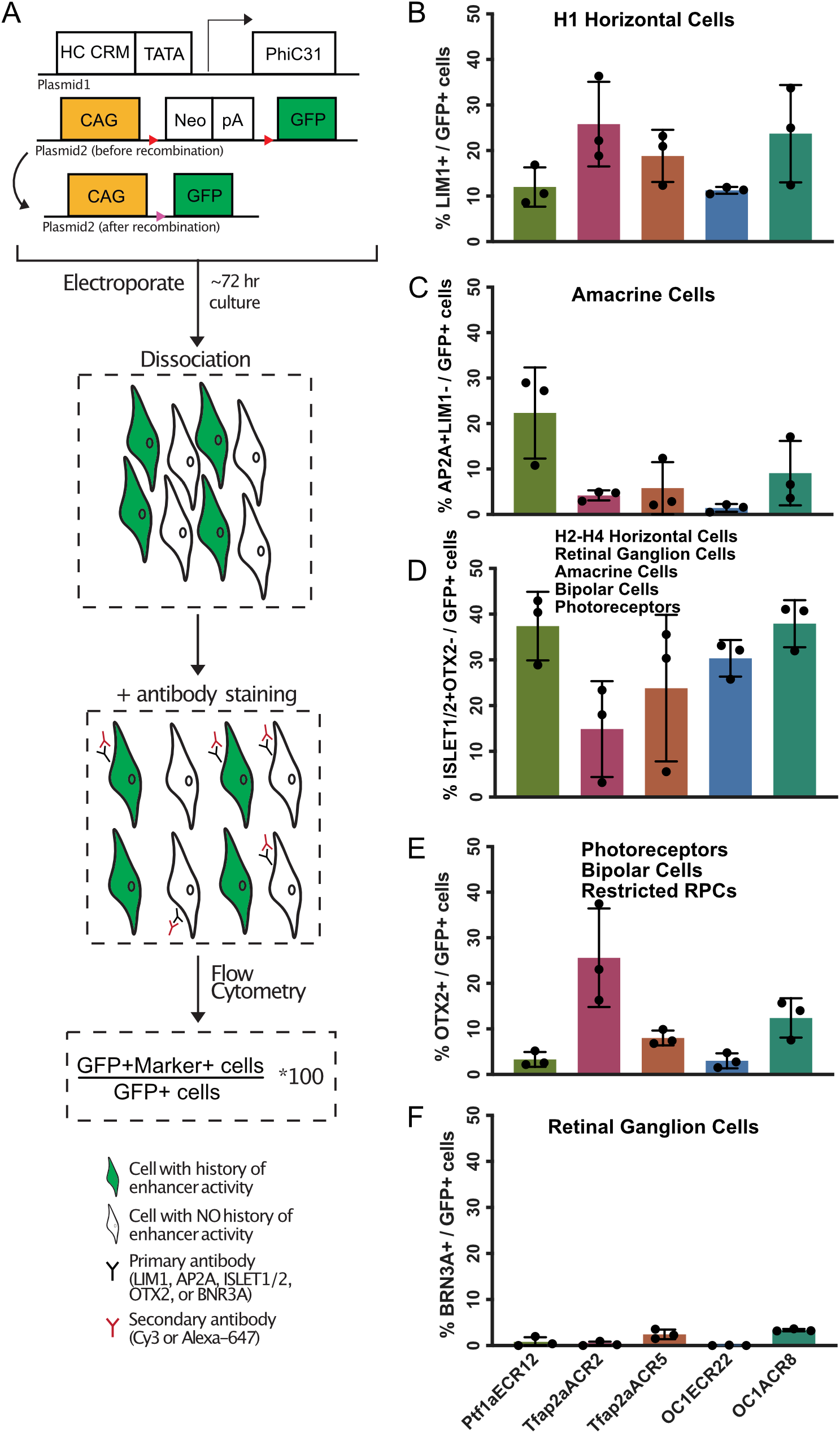
HC CRMs display varying levels of specificity to horizontal cells. A. Schematic of experimental workflow. B-F. Quantification of the percentage of cells with Ptf1aECR12, Tfap2aACR2, Tfap2aACR5, OC1ECR22 and OC1ACR8 activity that express various cell markers. Retinas were electroporated *ex vivo* at E5 with HC CRM::PhiC31 and CAaNa::GFP. After 3 days in culture, retinas were dissociated into single cells and counterstained for LIM1 and AP2A (B-C), ISLET1/2 and OTX2 (DE), or BRN3A (F) before being analyzed by flow cytometry. Error bars represent SEM. CRM, cis-regulatory module; pA, poly adenylation; RPC, retinal progenitor cell.

Next, cells were stained for ISLET1/2 and OTX2. ISLET1 is expressed in a variety of cell types including H2-H4 HCs, some ACs, bipolar cells, RGCs and photoreceptors, while ISLET2 marks a subset of photoreceptors and RGCs (Edqvist et al., 2006). OTX2 is expressed in photoreceptors and bipolar cells (Koike et al., 2007), and has also been shown to be expressed in at least two types of restricted retinal progenitor cells (Emerson et al., 2013; Ghinia Tegla et al., 2020). Therefore, cells that expressed ISLET1/2 alone were considered to be Islet1-expressing cells, identified as H2-H4 HCs, ACs or RGCs. The cells with expression of ISLET1/2 and OTX2, as well as those that only expressed OTX2, were identified as ISLET2-negative photoreceptors, bipolar cells or restricted progenitor cells. All 5 enhancers had activity in ISLET1 cells, which represent a range of 14-37% of cells with enhancer activity (Figure 3D). Only Tfap2aACR2 had substantial activity in OTX2 cells, which was expected based on the qualitative assessment of Tfap2aACR2 lineage tracing (Figure S3), while OTX2-positive cells represent 2-12% of the other enhancer populations (Figure 3E). Lastly, cells were stained for BRN3A which, in the mouse, marks 70% of retinal ganglion cells (Xiang et al., 1995). All enhancers had minimal activity in these RGCs, which represent a range of 0-3% of cells with enhancer activity (Figure 3F). These experiments were repeated and analyzed by confocal microscopy, yielding the same qualitative results (Figure S4). Upon these analyses, we have shown that Ptf1aECR12 is active almost entirely in HCs and ACs and that Tfap2aACR2 is largely active in HCs and cells expressing OTX2, while Tfap2aACR5 activity is more restricted and is primarily active in HCs. Most of OC1ACR8 activity is in HCs, although there is also activity in ACs and cells expressing OTX2, and finally, the activity of OC1ECR22 is nearly exclusive to HCs.

### PTF1A and RBPJ regulation of the OC1ECR22 element and OC1 expression

We sought to assess whether the activity of OC1ECR22 is consistent with its serving as an enhancer for OC1. In fact, 74.4% ±2.96 of cells with OC1ECR22 activity express OC1, as compared to only 18% ±1.56 of a control population (mean ± SEM, n=3, p<0.0001) (Figure 4A). This indicates that OC1ECR22 is likely an enhancer for the OC1 gene, and we were therefore interested in determining the transcription factors that activate the enhancer and thereby regulate OC1 expression. We performed motif analyses on the OC1ECR22 sequence with the MEME Suite to identify potential binding sites, and also searched manually through the sequence for additional relevant sites. Among the TF consensus sites that we identified were those for PTF1A and RBPJ (Figure 4B), two factors that have been shown to function as part of a trimeric complex along with an E-protein (Beres et al., 2006). These sites are separated by 5 nucleotides, which is in accordance with previously described PTF1A (E-Box) and RBPJ (TC-Box) binding sites separated by one helical turn of DNA (Masui et al., 2008). In addition, these factors have previously been shown to work together to regulate HC development (Lelièvre et al., 2011). To test the necessity of these sequences for cis-regulatory activity, we created versions of OC1ECR22 with mutations in either one or both of these sites and electroporated the plasmids into E5 chick retinas alongside the full version of OC1ECR22, and CAG::iRFP as an electroporation control. There was decreased activity of the enhancer upon mutation of the PTF1A site, with activity in 10.2% ±1.18 of electroporated cells compared to 29.8% ±4.24 in the control. Similarly, mutation of the RBPJ site led to activity in only 4.38% ±0.44 of electroporated cells (mean ± SEM, n=4, p<0.001) (Figure 4C-F). Finally, the joint mutation of the PTF1A and RBPJ sites resulted in the complete abrogation of OC1ECR22 activity, with activity in only 0.43% ±0.1 of electroporated cells compared to 25.6% ±2.24 (mean ± SEM, n=4, p<0.0001) (Figure 4G-I). Mutation of these sites, as well as mutation of a site that shows similarity to a nuclear hormone receptor binding site, are unlike multiple other sequence mutations that did not affect OC1ECR22 activity (Figure S5). Furthermore, overexpression of PTF1A and RBPJ resulted in a large increase in the activity of OC1ECR22, from 25.6% ±2.24 to 85.7% ±3.77 of electroporated cells, but no increase in the mutated version of OC1ECR22 (mean ± SEM, n=4, p<0.0001) (Figure 4G). We analyzed a previously reported ChIP-Seq dataset for Ptf1a in the mouse neural tube and detected enriched binding at a sequence that is the mouse homolog of OC1ECR22 (Figure S6) (Borromeo et al., 2014). This provides compelling evidence that PTF1A and RBPJ are binding to OC1ECR22 to regulate its activity.

**Figure 4.**
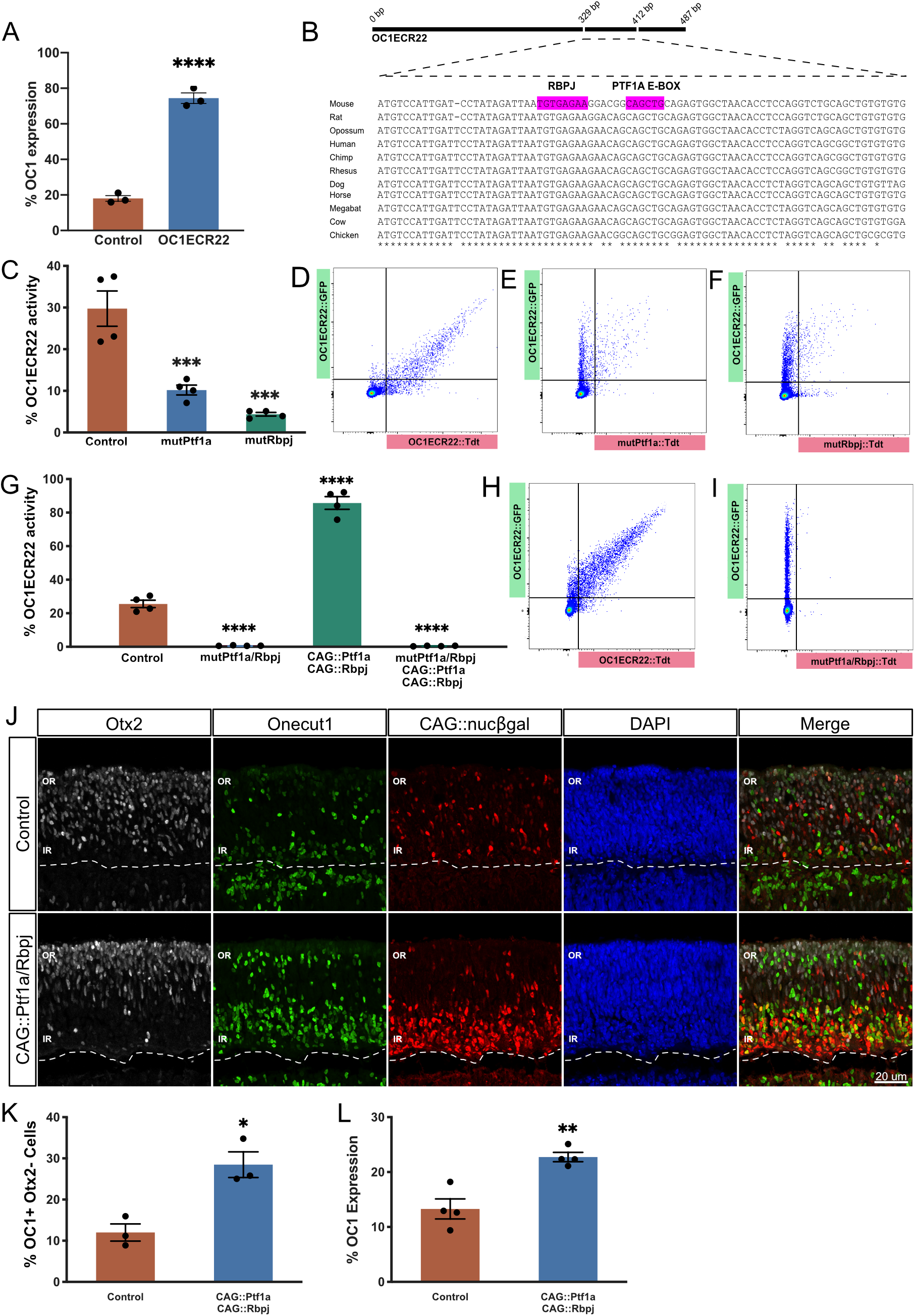
PTF1A and RBPJ regulate OC1 expression through binding and activation of OC1ECR22. A. Quantification of the percentage of control or OC1ECR22 cells that express OC1. Retinas were electroporated *ex vivo* at E5 with CAG::iRFP (control) and OC1ECR22::GFP. After two days in culture, retinas were dissociated and counterstained for OC1 before being analyzed by flow cytometry. Error bars represent SEM, n=3, p<0.0001 upon unpaired t test with two tailed distribution. B. View of OC1ECR22 sequence alignment with RBPJ and PTF1A E-box binding sites highlighted in magenta. Asterisks indicate conserved nucleotides, parenthesis contain consensus sites. C. Quantification of the percentage of electroporated cells with control or mutated OC1ECR22 activity. Retinas were electroporated *ex vivo* at E5 with CAG::iRFP as an electroporation control, OC1ECR22::GFP and either OC1ECR22::Tdt (control), mutPtf1a::Tdt, or mutRbpj::Tdt. After 2 days in culture, retinas were dissociated and analyzed by flow cytometry. Error bars represent SEM, n=4, p<0.001 upon one-way ANOVA with Dunnett’s multiple comparison test. D-F. Representative dot plots of retinas quantified in C. G. Quantification of the percentage of electroporated cells with control or mutated OC1ECR22 activity upon overexpression of PTF1A and RBPJ. Retinas were electroporated *ex vivo* at E5 with CAG::iRFP as an electroporation control, OC1ECR22::GFP, and OC1ECR22::Tdt (control) or mutPtf1a/Rbpj::Tdt, with or without CAG::Ptf1a and CAG::Rbpj. After two days in culture, retinas were dissociated and analyzed by flow cytometry. Error bars represent SEM, n=4, p<0.0001 upon one-way ANOVA with Dunnett’s multiple comparison test. H-I. Representative dot plots of retinas quantified in G. J. OTX2 and OC1 mark OC1+OTX2-horizontal cells. Representative images of retinas electroporated *ex vivo* at E5 with CAG::nucβgal as an electroporation control, with or without CAG::Ptf1a and CAG::Rbpj. After two days in culture, retinas were counterstained for Otx2 and OC1. Maximum intensity projection of 40x image, scale bars represent 20 μm. K. Quantification of the percentage of electroporated cells that are OC1+OTX2-upon overexpression of PTF1A and RBPJ as shown in J. Error bars represent SEM, n=3, p<0.05 upon unpaired t test with two tailed distribution. L. Quantification of the percentage of electroporated cells that express OC1 upon overexpression of PTF1A and RBPJ. Retinas were electroporated *ex vivo* at E5 with CAG::iRFP as an electroporation control with or without CAG::Ptf1a and CAG::Rbpj. After 2 days in culture, retinas were dissociated and counterstained for OC1 before being analyzed by flow cytometry. Error bars represent SEM, n=4, p<0.01 upon unpaired t test with two tailed distribution. ECR, evolutionary conserved region; OR, outer retina; IR, inner retina.

If OC1ECR22 serves as an enhancer element for OC1, the activation of OC1ECR22 by PTF1A and RBPJ could be translated to an effect on OC1 expression. We therefore overexpressed PTF1A and RBPJ in E5 retinas, alongside CAG::nβgal as an electroporation control. We observed that the joint overexpression of PTF1A and RBPJ resulted in a qualitative shift of the electroporated cells toward the inner retina, possibly indicating that more HCs were produced (Figure 4J). To quantify the OC1 expression among the electroporated cells in the inner retina, OTX2 was used as a counterstain. Previously, OC1 and OTX2 were shown to be expressed in a class of restricted retinal progenitor cells that gives rise to cone photoreceptors and HCs (Emerson et al., 2013). The OC1 cells that are not marked by OTX2 are therefore likely to be HCs, and those were the cells that we quantified. The percentage of OC1 expression in the electroporated cells of the inner retina doubled upon overexpression of PTF1A and RBPJ, from 12% ±2.08 to 28.5% ±3.12 of electroporated cells (mean ± SEM, n=3, p<0.05) (Figure 4K). Similarly, OC1 expression increased significantly from 13.3% ±1.82 to 22.7% ±0.85 upon overexpression of PTF1A and RBPJ when quantified by FACS (mean ± SEM, n=4, p<0.01) (Figure 4L). This, taken in combination with the effect of PTF1A and RBPJ on OC1ECR22 activation, suggests that PTF1A and RBPJ are direct activators of OC1 expression in developing HCs.

### Motif 4 is responsible for activation of the Tfap2aACR5 element

Tfap2aACR5 is another element that is predominantly active in HCs. Furthermore, 40.62% ±2.73 of cells with Tfap2aACR5 activity express AP2A, compared to only 11.1% ±0.7 of a control population (mean ± SEM, n=4, p<0.0001) (Figure 5B). This indicates that Tfap2aACR5 is likely an enhancer for the TFAP2A gene, and we were again interested in determining the factors that activate the enhancer. We created truncated versions of the enhancer so that the required portions of this 1.4 kb sequence could be identified. Five motifs of evolutionary conservation were identified within this sequence, and are likely to contain the sites required for the activity of Tfap2aACR5. We created a series of truncations within Tfap2aACR5 in the context of a GFP reporter plasmid and individually co-electroporated each one into E5 chick retinas with a fulllength Tfap2aACR5 TdTomato reporter and CAG::iRFP as an electroporation control, and analyzed them by flow cytometry after two days in culture. We first deleted a region that contained motifs 1, 2, and 3, and observed no change in Tfap2aACR5 activity. We next extended this deletion to include motif 4, and observed a significant decrease in reporter activity. Tfap2aACR5 was active in 10.3% ±0.31 of electroporated cells and the truncated version was active in only 0.86% ±0.23 of electroporated cells, indicating that motif 4 contains critical sequence elements for Tfap2aACR5 activity (mean ± SEM, n=3, p<0.001) (Figure 5A). A deletion of the remaining motif 5 did not lead to any further loss of Tfap2aACR5 activity.

**Figure 5.**
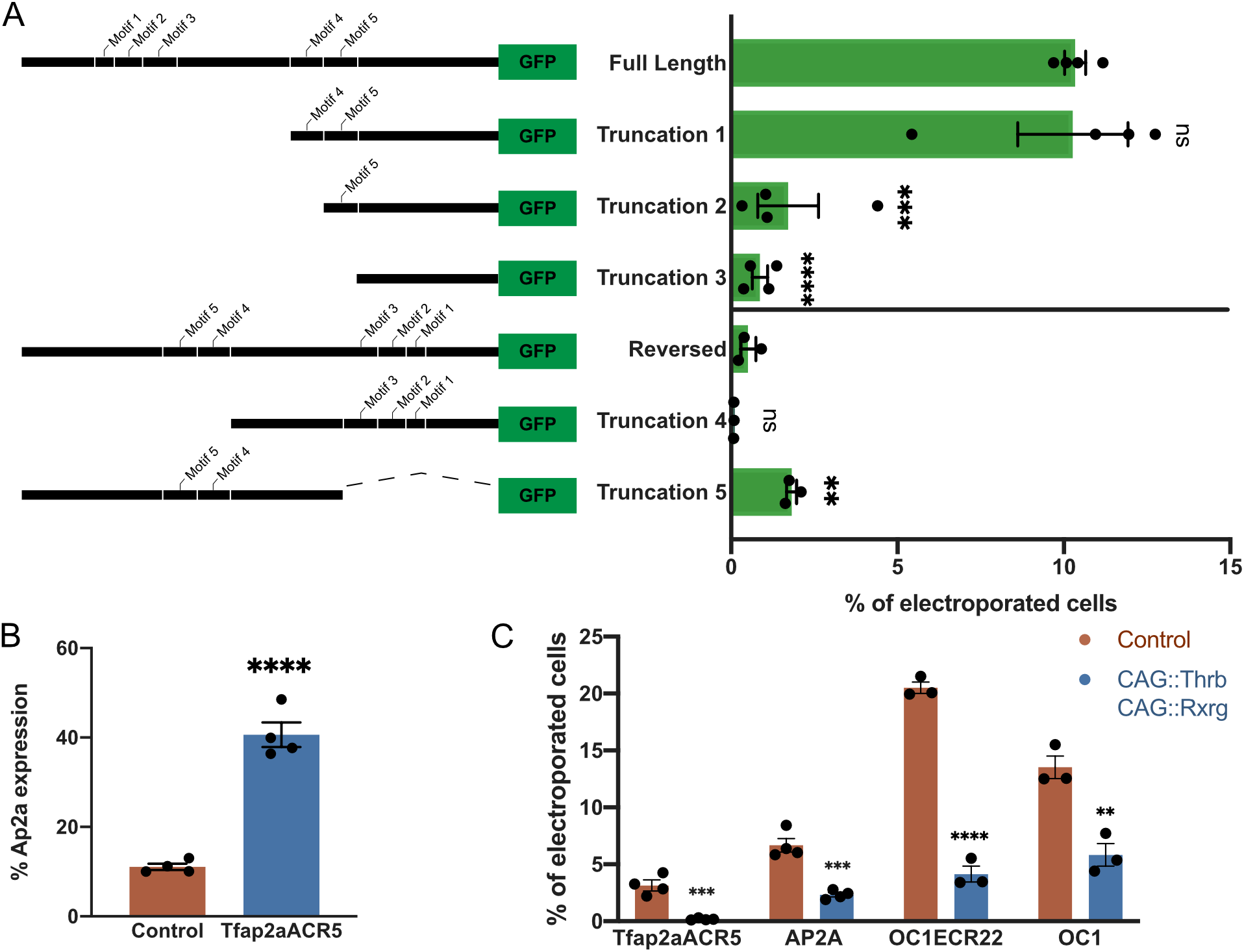
THRB and RXRG negatively regulate Tfap2aACR5 and OC1ECR22 activity and AP2A and OC1 expression. A. Truncations of the Tfap2aACR5 sequence reveal regions important for activity. Retinas were electroporated *ex vivo* at E5 with CAG::iRFP as an electroporation control, Tfap2aACR5::Tdt, and Tfap2aACR5::GFP or Tfap2aACR5trunc::GFP. After two days in culture, retinas were dissociated and analyzed by flow cytometry. Quantification of the percentage of electroporated cells with GFP driven by a truncated version of Tfap2aACR5. Error bars represent SEM, n=3 or 4, p<0.01 upon one-way ANOVAs with Dunnett’s multiple comparison test. B. Quantification of the percentage of control or Tfap2aACR5 cells with AP2A expression. Retinas were electroporated *ex vivo* at E5 with CAG::iRFP (control) and Tfap2aACR5::GFP. After 2 days in culture, retinas were dissociated and counterstained for AP2A before being analyzed by flow cytometry. Error bars represent SEM, n=4, p<0.0001 upon unpaired t test with two tailed distribution. C. Quantification of the percentage of electroporated cells with Tfap2aACR5 and OC1ECR22 activity and with AP2A and OC1 expression. Retinas were electroporated *ex vivo* at E5 with CAG::iRFP as an electroporation control, Tfap2aACR5::GFP or OC1ECR22::GFP, with or without CAG::Thrb and CAG::Rxrg. After 2 days in culture, retinas were dissociated and counterstained for AP2A or OC1 before being analyzed by flow cytometry. Error bars represent SEM, n=3 or 4, p<0.01 upon unpaired t tests with two tailed distribution. ACR, accessible chromatin region; ECR, evolutionary conserved region.

Although enhancers are thought to be active in both orientations, we have observed that some orientation-specific activity does occur. Tfap2aACR5 is one of the enhancers that has significantly decreased activity in the reverse orientation, with activity in only 0.51% ± 1.3 of electroporated cells compared to 4.96% ±1.52 of electroporated cells in the other orientation (mean ± SEM, n=3, p<0.05) (Figure 5A). Nevertheless, to test the requirement of motifs without altering the distance between remaining motifs and the basal promoter, we created two truncations of the reversed Tfap2aACR5. When a region that includes motifs 4 and 5 was deleted, there was again a decrease in the activity of the enhancer. When a region that includes motifs 1, 2, and 3 was deleted, we saw a significant increase in the activity of Tfap2aACR5, potentially indicating that repression mediated through those motifs was relieved. The reversed Tfap2aACR5 was active in 0.51% ± 1.3 of electroporated cells and the truncated version was active in 1.8% ± 0.004 of electroporated cells (mean ± SEM, n=3, p<0.01) (Figure 5A). These analyses identified a 100 bp region necessary for Tfap2aACR5 activation, and have introduced the possibility of repressors binding to negatively regulate enhancer activity.

### Nuclear receptors negatively regulate HC enhancer activity and gene expression

Among the sites identified upon the initial motif analysis of Tfap2aACR5 were half-sites that could be bound by several nuclear receptors, which include THRB, RXRG, ESRRG, and RORB. However, the mutation of these sites did not result in any change in the activity of the enhancer (Figure S7). In parallel, we overexpressed THRB and RXRG in combination with Tfap2aACR5 or OC1ECR22 in E5 chick retinas, which were then analyzed by flow cytometry after two days in culture. We observed significantly decreased activity of both enhancer elements, with Tfap2aACR5 activity decreasing from 3.14% ±0.48 to 0.19% ± 0.04 of electroporated cells, and OC1ECR22 activity decreasing from 20.51% ±0.5 to 4.14% ±0.7 of electroporated cells upon overexpression (mean ± SEM, n=3 or 4, p<0.001) (Figure 5C). Furthermore, when we stained the dissociated cells for AP2A and for OC1 we observed similar decreases. The overexpression of THRB and RXRG resulted in 2.32% ±0.19 of cells expressing AP2A, compared to 6.67% ±0.59 of cells in the control condition, and 5.83% ±0.99 of cells expressing OC1 compared to 13.52% ±0.99 of cells in the control condition (mean ± SEM, n=3 or 4, p<0.01) (Figure 5C). This decrease in AP2A and OC1 expression is presumably mediated by the decreased enhancer activity. These data introduced the possibility that these nuclear receptors are serving as negative regulators of HC genes in developing photoreceptors, although this interaction could be indirect given that the identified half-sites in Tfap2aACR5 were not required for this effect.

## Discussion

Identification of cis-regulatory elements serves an important role in the study of gene regulatory events that drive the formation of specific cell types. By surveying the genomic regions surrounding several genes known to be functionally relevant to HC development of and screening the DNA elements for specific activation, we have identified OC1ECR22 and Tfap2aACR5 as novel enhancer elements with preferential activity in HCs. There is a twofold use to these elements. Firstly, there is a notable lack of DNA elements that can be used to label HCs during development, which would allow for the monitoring and isolation of those cells. Secondly, these elements represent nodes of the gene regulatory networks that underlie HC development, and as such, can be used to identify the connections between nodes. Although there are several known enhancer elements for genes associated with HC populations, such as PTF1A and OC1, there has not been extensive validation of their activity in HCs. Both a 2.3 kb regulatory region upstream and a 12.4 kb regulatory region downstream of PTF1A drive expression in cells of the retina, and while the 5’ enhancer has been shown to function as part of an autoregulatory loop dependent on PTF1A and RBPJ expression, the 3’ enhancer contains a 0.8 kb sub-fragment that is responsive to FOXN4 and RORB (Liu et al., 2013; Masui et al., 2008; Meredith et al., 2009). The inactive Ptf1aECR2 element described in our study aligns to the 5’ enhancer, and several CRMs align to the 3’ enhancer. These include the Ptf1aECR12, Ptf1aECR27 and Ptf1aACR4 elements which are active in the retina, and the Ptf1aECR7, Ptf1aECR9, Ptf1aECR10, Ptf1aECR11, Ptf1aECR13, and Ptf1aECR14 elements which are not active in the retina. Additionally, our lab has recently reported a screen for regulatory elements that direct OC1 expression in restricted RPCs which generate HCs (Patoori et al., 2020). Here, we characterized a subset of cis-regulatory elements identified in this screen that are preferentially active in HCs (OC1ECR22, OC1ACR4 and OC1ACR8) which suggests that there is a distinct gene regulatory network that is activated to increase OC1 expression in newly formed HCs.

In this study, we have utilized recombinase-based lineage tracing analysis to ensure that information relating to enhancer activity is not lost due to their transient activation during development. In fact, we have shown that a history of activity in entirely new populations of cells can be revealed upon lineage tracing, as is the case with marked photoreceptors observed upon lineage tracing of the Tfap2aACR2 element. In contrast, only a minimal percentage of Tfap2aACR2::GFP labeled populations express OTX2 (data not shown). Furthermore, several of the previously reported enhancer elements that are active in HCs are also active in other early-born cell types such as photoreceptors and retinal ganglion cells. These include the ThrbCRM1, ThrbICR and Rxrg208 elements (Blixt and Hallböök, 2016; Emerson et al., 2013; Jones et al., 2007). Therefore, the strict screening measures described above are crucial to ensure that these novel elements can be used to specifically mark developing HCs. Interestingly, we found that several elements drove expression of reporters in cell types that are unexpected when considering the genes that they are located near. This was the case for elements selected near all four targeted genes and is especially striking in the case of LIM1 which is exclusively expressed in HCs. It is of course possible that although the DNA elements in question were selected based on proximity to PTF1A, OC1, TFAP2A or LHX1, they may not be involved in the transcriptional regulation of those genes but of others that are more distally located. Another potential explanation is that there are several elements for each gene that must be active in combination to lead to transcriptional regulation in specific cell types, so that the pattern of activity driven by an individual enhancer does not accurately reflect the ultimate pattern of gene expression.

Motif analysis of the OC1ECR22 element enabled us to determine that PTF1A and RBPJ act together to activate OC1ECR22 and thereby regulate OC1 expression. Subsequently, we detected enriched binding of PTF1A at the OC1ECR22 sequence upon analysis of ChIP-Seq data from the mouse neural tube (Borromeo et al., 2014). However, we did not detect enriched binding of RBPJ at this site, which could indicate that RBPJ is not present but could also represent a technical limitation of detection of RBPJ. In addition to supporting our finding that PTF1A directly promotes OC1 expression in HCs via the OC1ECR22 element, this suggests that there may be a regulatory relationship between PTF1A and OC1 in other portions of the nervous system. While both PTF1A and OC1 have been shown to be essential for HC fate determination, a regulatory relationship between the two has not been reported (Fujitani et al., 2006; Wu et al., 2013). This conclusion was reached as there was no change in PTF1A expression observed in OC1-null retinas and no change in OC1 expression observed in PTF1A-null retinas, although the small number of cells that co-expressed the two factors may have impacted that analysis (Wu et al., 2013). In the same study, overexpression of PTF1A did not result in increased expression of OC1. We observed that co-overexpression of PTF1A and RBPJ led to increased OC1 expression and to cell body localization changes in the electroporated population that indicate increased HC genesis. However, it is possible that the individual overexpression of PTF1A or RBPJ are not able to induce these changes. If that is the case, it would suggest that endogenous RBPJ is limiting with regards to PTF1A overexpression and could underlie the discrepancy in the conclusions of this study and Wu et al., 2013.

We propose a model for the transcriptional regulation of select HC genes, as well as for the relationships between this cis-regulatory activity and specific cell types. FOXN4 and RORB are critical for HC development, and have been shown to regulate PTF1A expression (Fujitani et al., 2006; Liu et al., 2013). Our data suggests that PTF1A, in combination with RBPJ, is sufficient to induce OC1 expression through the OC1ECR22 element. Additionally, increased levels of THRB and RXRG expression results in the negative regulation of OC1 and AP2A expression mediated through the OC1ECR22 and Tfap2aACR5 elements (Figure 6A). While this study focused on the interaction between PTF1A and OC1, the cis-regulatory elements that have been reported here can provide a foothold to examine the relationships between additional factors in the HC gene regulatory network. The HC CRMS for PTF1A, OC1 and TFAP2A that were extensively characterized in this study display activity in cone photoreceptors, HCs, ACs and RGCs. These cell types, with the exclusion of ACs, have previously been reported to be derived, at least in part, from restricted RPCs (rRPCs) that are in turn born from multipotent RPCs (mRPCs) (Emerson et al., 2013; Hafler et al., 2012; Jones et al., 2007; Suzuki et al., 2013). The Ptf1aECR12 element is active in developing HCs and ACs (Figure 6B), which is consistent with PTF1A expression in those cells. The OC1ACR8 element is active in cone photoreceptors in addition to HCs and ACs, while OC1ECR22 activity is more restricted to HCs (Figure 6C). We did however detect some RGCs marked by the OC1ECR22 element through *in vivo* lineage tracing performed at E3 (data not shown). This suggests that OC1ECR22 is active in the RGC population prior to the onset of the *ex vivo* electroporation experiments initiated at E5. Finally, the Tfap2aACR2 element is active in cone photoreceptors and HCs, while Tfap2aACR5 is more restricted to HCs (Figure 6D).

**Figure 6.**
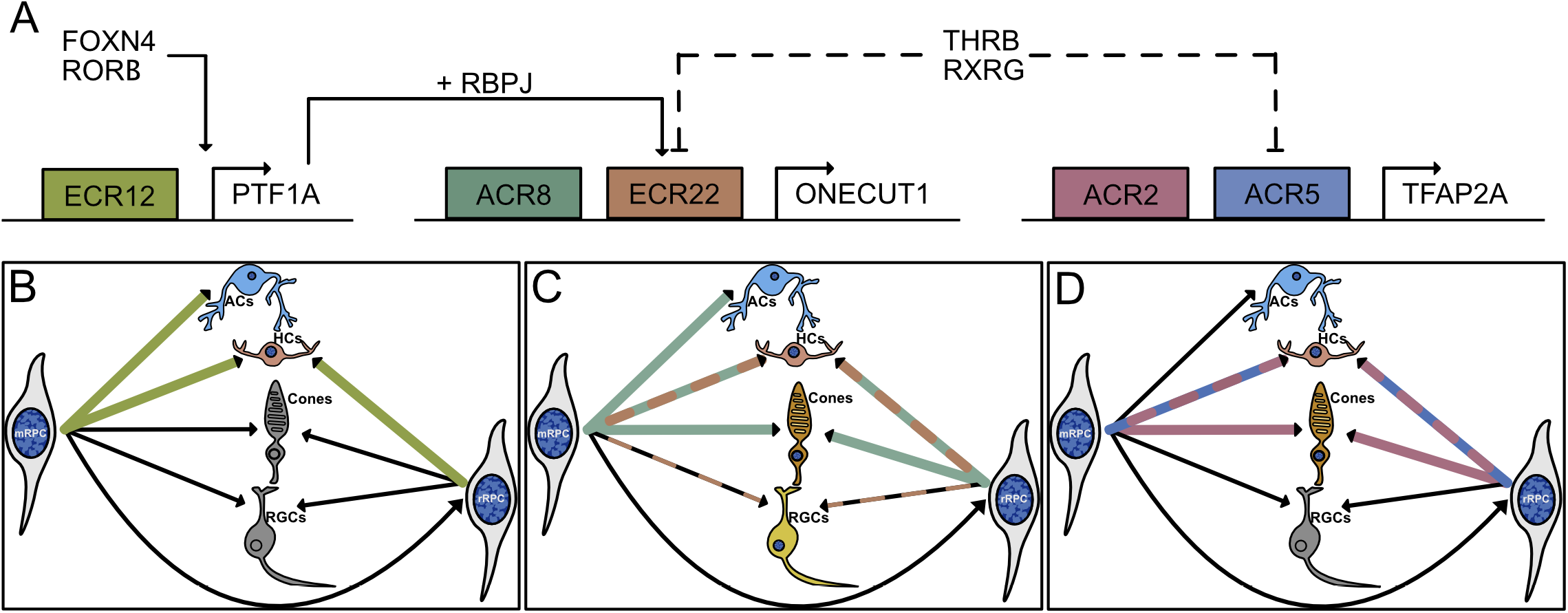
A proposed model of cis-regulatory activity during horizontal cell development. A. PTF1A, which is regulated by FOXN4 and RORB, acts in combination with RBPJ to regulate OC1 through activation of the OC1ECR22 element. THRB and RXRG may act indirectly to negatively regulate OC1 and AP2A expression via the OC1ECR22 and Tfap2aACR5 elements. B-D. Cell populations marked by activity of Ptf1aECR12 (B), OC1ACR4 and OC1ECR22 (C), and Tfap2aACR2 and Tfap2aACR5 (D). mRPC, multipotent retinal progenitor cell; rRPC, restricted retinal progenitor cell; AC, amacrine cell; HC, horizontal cell; RGC, retinal ganglion cell.

While PTF1A and RBPJ regulate OC1 transcription in HCs, they are likely not the only gene regulatory network input. In fact, we also observed that the deletion of a site within the OC1ECR22 sequence, with some similarity to a nuclear hormone receptor binding site, results in the complete loss of enhancer activity. This could serve as a binding site for RORB, a known regulator of HC development (Liu et al., 2013). This highlights the complex nature of transcriptional control of gene regulation during development, and illustrates that there are many more details of the gene regulatory networks active in HCs to be uncovered. One of these steps is the activation of TFAP2A. We have identified a potential cis-regulatory element in the form of Tfap2aACR5 and identified a critical 100 bp activation region. However, we have not yet determined the sites in this region that are required for enhancer activity. Further testing will therefore be necessary to investigate and identify the factors responsible for regulating AP2A expression in developing HCs.

Overexpression of THRB and RXRG resulted in decreased HC enhancer activity as well as decreased expression of AP2A and OC1, introducing the possibility that these factors are functioning in restricted progenitor cells or in developing photoreceptors to negatively regulate HC gene expression. It is possible that this would be mediated indirectly, as the three steroid hormone receptor binding sites within the Tfap2aACR5 sequence were dispensable to its regulation. We were interested in observing the loss-of-function phenotype for THRB and RXRG, and performed CRISPR/Cas9 gene editing of these genes via electroporation (data not shown). We did not detect any changes in the photoreceptor or HC populations which may indicate that the effects observed with ectopic expression of THRB and RXRG does not reflect their *in vivo* function, but could also be due to a number of technical limitations. For example, we are currently unable to validate loss of THRB and RXRG protein in cells using antibodies and therefore cannot determine the efficacy of our gene editing strategy to create loss-of-function alleles. Another possibility is that THRB and RXRG are not directly involved in HC gene regulation, but their overexpression interferes with the normal function of a nuclear receptor, such as RORB, that is critical for HC differentiation. Further experiments will be necessary to clarify the role of THRB and RXRG in relation to HC gene expression during retinal development.

## Methods

### Animals

Fertilized chicken eggs were obtained from Charles River and were stored at 16 °C for 0-10 days. The eggs were then incubated in a 38 °C humidified incubator for 5 days.

### Bioinformatic Analysis

Candidate CRMs near PTF1A were identified based on conservation criteria using the ECR Browser and the UCSC Genome Browser programs (Kent et al., 2002; Ovcharenko et al., 2004). Candidate CRMs near PTF1A, TFAP2A, LHX1 and ONECUT1 were identified based on open chromatin profiles upon analysis of ATAC-Seq data aligned to the chicken genome (galGal5) in the UCSC Genome Browser (Patoori et al., 2020). To identify TF binding sites, sequence homologs of OC1ECR22 and Tfap2aACR5 were compiled using the UCSC Genome Browser, and were then aligned with the ClustalW program version 2.1 (Thompson et al., 1994). Conserved motifs within these CRMs were identified using MEME version 4.12.0 with the number and width of motifs adjusted to cover the length of the sequence (Bailey and Elkan, 1994). The MEME text output was then run through Tomtom version 4.12.0 where binding sites were predicted (Gupta et al., 2007). Transcription factor motifs were identified using the Eukaryote DNA, vertebrates (in vivo and in silico) database, which searches the JASPAR2018 CORE vertebrates, Uniprobe Mouse, and Jolma2013 Human and Mouse databases.

### DNA Plasmids

Stagia3 (Billings et al., 2010), CAaNa::GFP, CAG::TdTomato (Schick et al., 2019), CAG::AU1 (Emerson and Cepko, 2011), CAG::GFP (Matsuda and Cepko, 2007) and CAG::iRFP (Buenaventura et al., 2018) were described previously and CAG::mCherry and CAG::nucβgal (Cepko Lab, Harvard Medical School) were obtained from the Cepko lab. To clone candidate CRMs into Stagia3, sequences were first PCR-amplified from mouse or chicken genomic DNA. The amplicons were cloned into pGEM-T Easy (Promega, A1360) and sub-cloned into Stagia3 with EcoR1, or with Sal1, Xho1 or Mfe1 if the CRM sequence included an Ecor1 site. OC1ECR22, OC1ACR4 and OC1ACR8 sequences were described previously (Patoori et al., 2020). OC1ECR22 and Tfap2aACR5 were also cloned into Statia (Jean-Charles et al., 2018) with EcoR1. To clone the CRMs into lineage tracing plasmids, digested CRMs were cloned into bp::PhiC31 (Schick et al., 2019) with EcoR1. To clone CAG::Thrb and CAG::Rxrg, the respective coding sequences were PCR-amplified from chicken cDNA with primers that included Age1 and Nhe1 restriction enzyme sites. The amplicons were digested and cloned into a pCAG-MCS vector digested with Age1 and Nhe1. The pCAG-MCS vector was cloned by modifying a pCAG vector with annealed oligos so that Age1 and Nhe1 sites are flanked by Ecor1 sites. To clone CAG::Ptf1a and CAG::Rbpj, mouse cDNA clones of Ptf1a (BC132505) and of Rbpj (BC051387) were obtained from Transomic Technologies. The PTF1A and RBPJ coding sequences were cloned into a pCAG vector with Ecor1 or Ecor1 and Not1, respectively. All of the plasmids described above were verified by restriction digestion and Sanger sequencing.

### Mutagenesis and Truncation Assays

To create mutations within the OC1ECR22 and Tfap2aACR5 sequences, scrambled DNA sequences were first generated using the Shuffle DNA tool from the Sequence Manipulation Suite (Stothard, 2000). These sequences were incorporated into a set of primers that spanned the potential TF binding sites to create mismatches. A second set of primers included the outermost portions of the CRM sequence as well as restriction sites to facilitate cloning. The mutations were created using overlap extension PCR with the initial CRM plasmid serving as a template, and the mutated sequences were cloned into Stagia3 or Statia plasmids.

To create truncated versions of Tfap2aACR5, primers were designed so that a portion of the sequence would be excluded during amplification. The truncated sequences were PCR-amplified using the initial CRM plasmid as a template, and the amplicons were digested using Ecor1 and Sal1 whose sites were appended to the primer sequences. The truncations were then cloned into Stagia3. All of the mutations and truncations of CRM sequences were verified by restriction digestion and sequencing. Mutations and truncations were tested by electroporation of the mutated/truncated CRM driving one reporter with the original version driving a second reporter.

### DNA Electroporation and Culture

All DNA electroporations were carried out as described previously (Emerson and Cepko, 2011) using a Nepagene NEPA21 Type II Super Electroporator. Briefly, E5 chicken retinas were dissected in 1:1 DMEM/F12 (Gibco, 11320082) and electroporated with a DNA plasmids using 5 pulses of 25 V with a 50 millisecond pulse length and 950 millisecond interpulse interval. DNA mixes contained 100 ng/μl of all plasmids with CAG promoters or with PhiC31, and 160 ng/μl of all other plasmids. Electroporated retinas were cultured for 2-3 days as described previously (Emerson et al., 2013).

### Retina Dissociation and Flow Cytometry

Any remaining retinal pigment epithelium and the condensed vitreous were dissected from cultured retinas in HBSS (Gibco, 14170112), which were then dissociated using a papain-based protocol as described previously (Worthington, L5003126) (Matsuda and Cepko, 2004). Retinal cells were subsequently fixed in 4% paraformaldehyde/1X PBS for 15 minutes at room temperature and washed 3X in 1X PBS, or if the cells were being prepared for immunohistochemistry, in 1X PBS with 0.1% Tween-20 (VWR, 97062-332). Cells were then filtered into 4 mL FACS tubes (BD Falcon, 352054) through 40 μm cell strainers (Biologix, 15-1040).

Dissociated cells were analyzed with a BD LSR II flow cytometer using 488, 561 and 633 lasers. All experiments included control retinas that were either non-electroporated, or electroporated with CAG::GFP, CAG::Tdt, or CAG::iRFP. These retinas were used to generate compensation controls and to define gates on cell plots. Compensation controls for experiments that involved immunohistochemistry will be discussed below. All flow cytometry data was analyzed with FlowJo software Version 10.2.

### Immunohistochemistry

Cultured retinas were fixed in 4% paraformaldehyde/1X PBS for 30 minutes, washed 3X in 1X PBS, and cryo-protected in 30% sucrose/0.5X PBS. Retinas were flash-frozen in OCT (Sakura Tissue-Tek, 4583) and sectioned with a Leica cryostat into 20 μm vertical sections that were collected on slides (FisherBrand, 12-550-15). All immunofluorescence staining of retinal sections was performed as described previously (Emerson and Cepko, 2011). A 1 μg/μl DAPI solution (VWR, TCA2412-5MG) was applied to the slides for nuclear staining prior to mounting in Fluoromount-G (Southern Biotech, 0100-01) with 34 x 60 mm coverslips (VWR, 48393-106).

Staining of dissociated retinal cells was carried out as follows: After the final wash in 1X PBS with 0.1% Tween-20 (PBT) following fixation, cells were filtered into 4 mL FACS tubes as described above and resuspended in PBT with 5% serum. The cells were blocked for 1 hour shaking at room temperature. During that time a concentrated antibody solution was created in PBT with 5% serum to be diluted in the block already on the cells. The primary antibody solution was added to the cells, which were then incubated overnight shaking at 4 °C. On the following day, the primary antibody solution was washed from the cells by adding 3 mL PBT. Cells were blocked in PBT with 5% serum for 30 minutes. A concentrated secondary antibody solution was diluted in the block already on the cells, and the cells were then incubated in the dark for 1 hour at room temperature. The primary antibodies were washed from the cells with 3 mL 1X PBS, and the cells were resuspended in 1X PBS prior to analysis by flow cytometry. Retinas that were prepared as compensation controls were treated as follows: Nonelectroporated retinas were not treated with any primary antibodies, but were treated with secondary antibodies. Single-stain controls were treated with individual sets of primary and secondary antibodies.

The primary antibodies used were: chicken anti-GFP (ab13970, Abcam, 1:2000), rabbit anti-GFP (A-6455, Invitrogen, 1:1000), chicken anti-β-galactosidase (ab9361, Abcam, 1:1000), mouse IgG1 anti-β-galactosidase (40-1a-s, DSHB, 1:20), mouse IgG1 anti-Visinin (7G4-s, DSHB, 1:500), mouse IgG1 anti-Lim1 (4F2-c, DSHB, 1:30), mouse IgG2b anti-Ap2a (3B5, DSHB, 1:200), mouse IgG2b anti-Islet1+2 (39.4D5, DSHB, 1:10), mouse IgG1 anti-Brn3a (MAB1585, EMD Millipore, 1:800), goat anti-Otx2 (Af1979, R&D Systems,1:400), and mouse IgG2b anti-HNF-6 (sc-376308, Santa Cruz, 1:200). All secondary antibodies were obtained from Jackson Immunoresearch and were appropriate for multiple labeling. Alexa 488- and 647-conjugated secondary antibodies were used at 1:400 and cy3-conjugated secondary antibodies were used at 1:250.

### Alkaline Phosphatase staining

After fixation, retinas were incubated with 1 mL of NTM buffer (100 mM NaCl, 100 mM Tris pH 9.5, 50 mM MgCl2) shaking at room temperature. After 15 minutes, the NTM was replaced with 1 mL NTM with 0.25 mg/mL NBT (VWR, 97061-412) and 0.125 mg/mL BCIP. The retinas were incubated in the dark with these substrates, shaking for 2-3 hours. A positive control was used to determine that the development of the AP stain was complete.

### Imaging and Image Processing

Images of AP stained retinas were acquired with a Zeiss AxioZoom V16 microscope using a PlanNeoFluarZ1x objective. All confocal images of vertically sectioned retinas were acquired with a Zeiss LSM710 inverted confocal microscope and ZEN Black 2015 21 Sp2 software. Images were acquired at 1024 x 1024 resolution with an EC Plan-Neofluar 40X/1.30 Oil DIC M27 objective. Images were analyzed with FIJI (Schindelin et al., 2012). Cells were counted using the Cell Counter plugin for ImageJ in Z-stacks from a minimum of three retinas in each condition. All figures were assembled using the Affinity Designer vector editing program, and any adjustments to brightness and contrast were applied uniformly across images. Schematics were assembled with Adobe Illustrator.

### Statistical Analysis

Graphs were made using GraphPad Prism8 software version 8.4.2, and error bars represent standard error of the mean (SEM). Statistical analyses were also performed using GraphPad Prism 8 software. Data was first tested for normality using the Shapiro-Wilk test. When data was found to be distributed normally, an unpaired t-test with two tailed distribution or one-way ANOVA with Dunnett’s multiple comparison test was performed, as noted.

## Acknowledgements

Support was provided by National Science Foundation grant CAREER 1453044 (to MME), National Eye Institute grant R01EY024982 (to MME) and NIMHD 3G12MD007603-30S2 (CCNY). The content is solely the responsibility of the authors and does not necessarily represent the official views of the National Institute On Minority Health and Health Disparities or the National Institutes of Health or the National Science Foundation. Jeffrey Walker and Jorge Morales provided excellent technical support with flow cytometry and confocal microscopy experiments. The 40-1a, 7G4, 4F2, 39.4D5, and 3B5 antibodies developed by Joshua Sanes, Suzanne Bruhn and Constance Cepko, Susan Brenner-Morton and Thomas Jessell, and Trevor Williams were obtained from the Developmental Studies Hybridoma Bank developed under the auspices of the NICHD and maintained by the Department of Biology at the University of Iowa. We thank Diego Buenaventura for the construction of the CAG::Thrb and CAG::Rxrg plasmids, Twinkle Jacob for the construction of the Ptf1aACR3 and Lhx1ACR4 plasmids, Stephen Madamba for the construction of the Ptf1aECR22::GFP plasmid, Emily Lamarre for the construction of the mutSite1::Tdt and mutSite2::Tdt plasmids, and the members of the Emerson lab for support throughout the project.

