## Supplementary material for "Early cis-regulatory events in the formation of retinal horizontal cells": Supp. Fig Legends

### Supplemental Figure Legends

**Figure S1.** AP reporter assay for HC CRM activity. Representative images of whole retinas electroporated *ex vivo* at E5 with CAG::mCherry as an electroporation control and HC CRM::PLAP. After 2 days in culture, AP stain was developed. Scale bars represent 200 µm.

**Figure S2.** Visinin counterstain rules out HC CRMs with activity in photoreceptors. Representative images of vertically sectioned retinas that were electroporated *ex vivo* at E5 with CAG::AU1 as an electroporation control and HC CRM::GFP. After 2 days in culture, retinas were stained for Visinin. Maximum intensity projection of 40x image, scale bars represent 20 µm.

**Figure S3.** Lineage tracing of active CRMs.
A-B. Representative images of vertically sectioned retinas that were electroporated *ex vivo* at E5 with CAG::nucβgal as an electroporation control, HC CRM::PhiC31 and CAaNa::GFP. After two days in culture, retinas were counterstained for Visinin to mark photoreceptors (A) or for LIM1 to mark horizontal cells (B). Maximum intensity projection of 40x image, scale bars represent 20 µm.

**Figure S4.** HC CRMs display varying levels of specificity to horizontal cells.
A-C. Representative images of vertically sectioned retinas that were electroporated *ex vivo* at E5 with CAG::nucβgal as an electroporation control, HC CRM::PhiC31 and CAaNa::GFP. After 3 days in culture, retinas were counterstained for LIM1 and AP2A (A), Visinin and ISLET1/2 (B), or OTX2 and BRN3A (C). Maximum intensity projection of 40x image, scale bars represent 20 µm.

**Figure S5.** Additional mutations in the OC1ECR22 sequence.
A. View of OC1ECR22 sequence alignment with OC1, OTX2, RBPJ, PTF1A E-box, and PTF1A E-box#3 binding sites highlighted in magenta. Asterisks indicate conserved nucleotides.
B-C. Quantification of the percentage of electroporated cells with control or mutated OC1ECR22 activity. Retinas were electroporated *ex vivo* at E5 with CAG::iRFP as an electroporation control, OC1ECR22::GFP and either OC1ECR22::Tdt (control), mutOtx2::Tdt (B), mutOC1::Tdt or mutPtf1a#3::Tdt (C). After 2 days in culture, retinas were dissociated and analyzed by flow cytometry. Error bars represent SEM, n=4.
D. View of OC1ECR22 sequence alignment with Site1 sequence underlined and Site2 sequence highlighted in gray. Asterisks indicate conserved nucleotides.
E. Quantification of the percentage of electroporated cells with control or mutated OC1ECR22 activity. Retinas were electroporated *ex vivo* at E5 with CAG::iRFP as an electroporation control, OC1ECR22::GFP and either OC1ECR22::Tdt (control) mutSite1::Tdt or mutSite2::Tdt. After 2 days in culture, retinas were dissociated and analyzed by flow cytometry. Error bars represent SEM, n=4, $p<0.001$ upon one-way ANOVA with Dunnett's multiple comparison test.

**Figure S6.** PTF1A binds to OC1ECR22 sequence.
A. View of PTF1A and RBPJ ChIP-Seq datasets from the spinal cord (Borromeo et al., 2014) aligned to the mm9 genome in UCSC Genome Browser at the OC1 locus.

B. Zoomed view of area enclosed in rectangle in A. Portion of OC1ECR22 corresponding to RBPJ and PTF1A sites enclosed in black rectangle.

**Figure S7.** Additional mutations in the Tfap2aACR5 element.
A. View of Tfap2aACR5 sequence alignment with THRB, RXRG, ESRRG, bHLH binding sites highlighted. Asterisks indicate conserved nucleotides.
B-E. Quantification of the percentage of electroporated cells with control or mutated Tfap2aACR5 activity. Retinas were electroporated *ex vivo* at E5 with CAG::iRFP as an electroporation control, OC1ECR22::GFP and either OC1ECR22::Tdt (control), mutEsrrg::Tdt or mutThrb::Tdt (B), mutRxrg::Tdt (C) mutThrb/Rxrg/Esrrg::Tdt (D), or mutbHLH::Tdt (E). After 2 days in culture, retinas were dissociated and analyzed by flow cytometry. Error bars represent SEM, n=4.
