## Supplementary figures and images for "Early cis-regulatory events in the formation of retinal horizontal cells"

### Figure S1

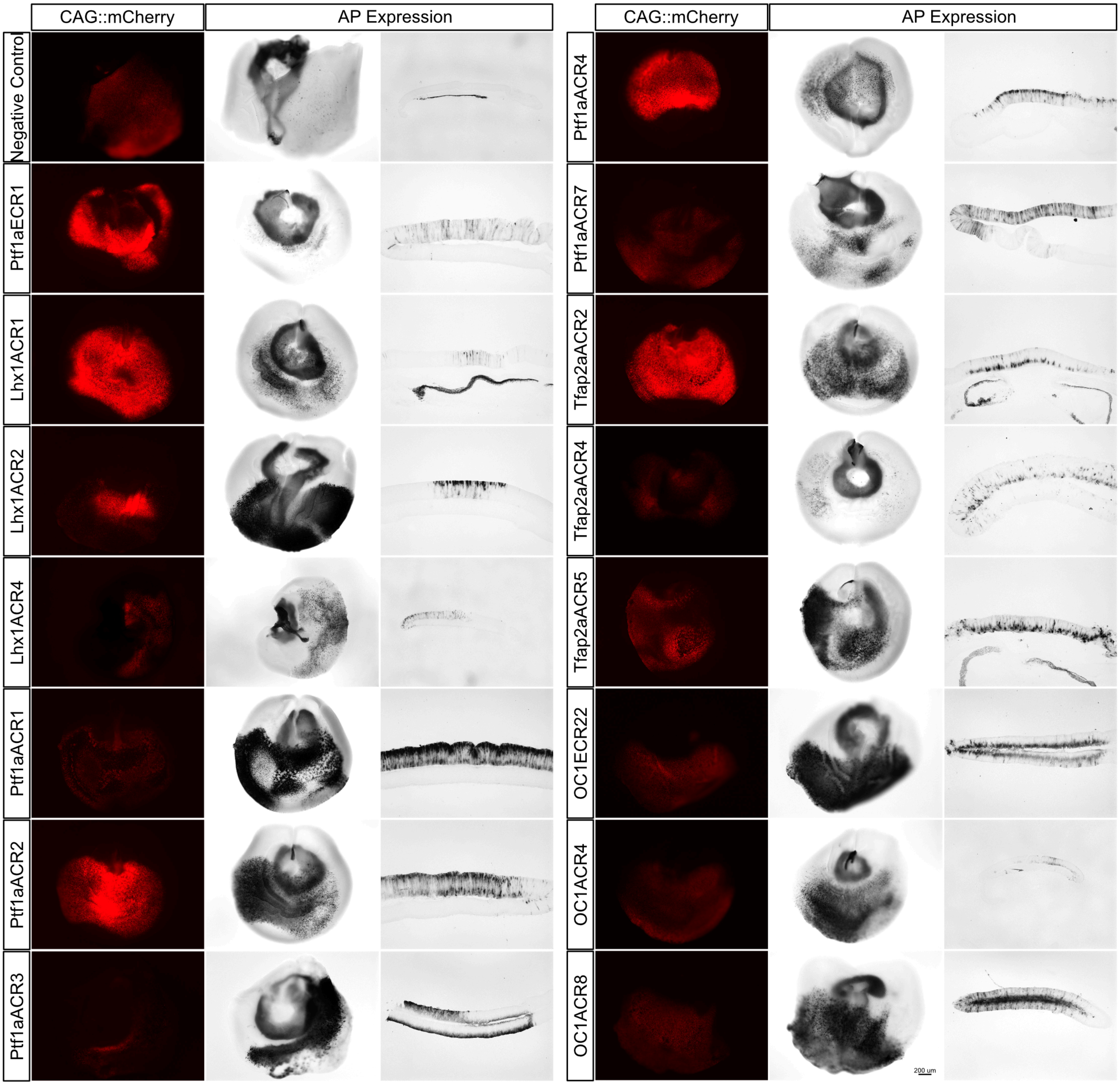

### Figure S2

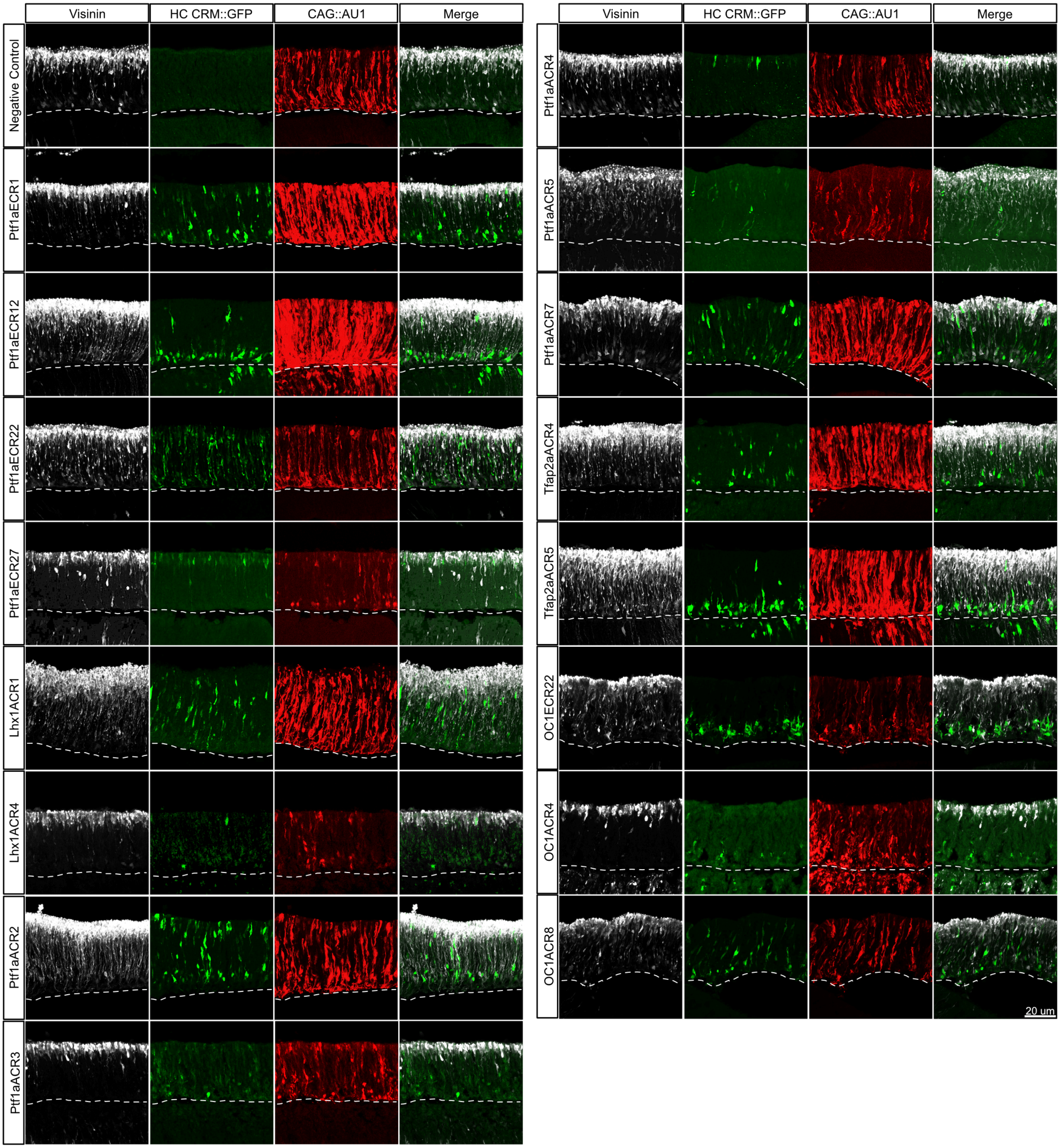

### Figure S3

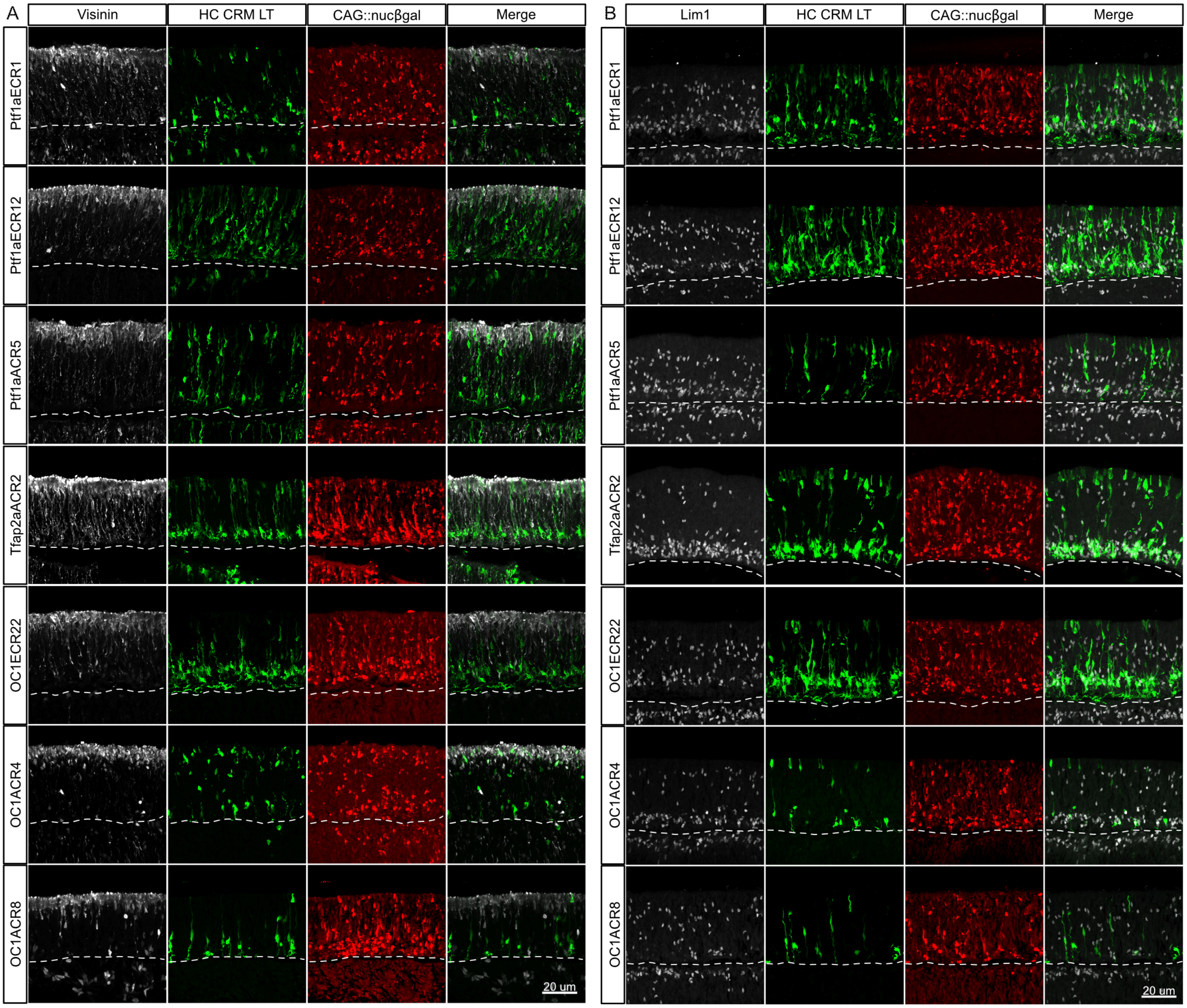

### Figure S4

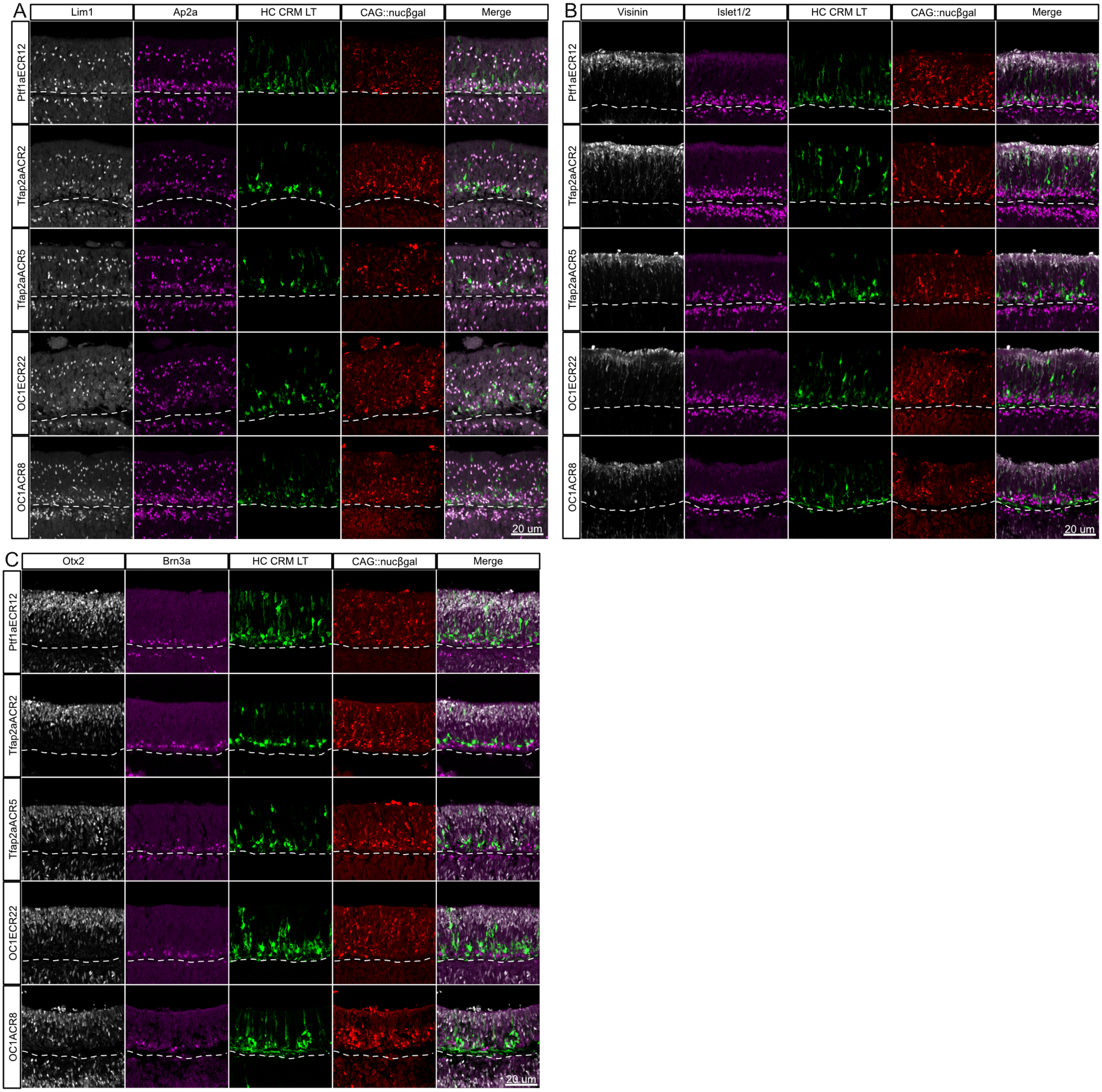

### Figure S5

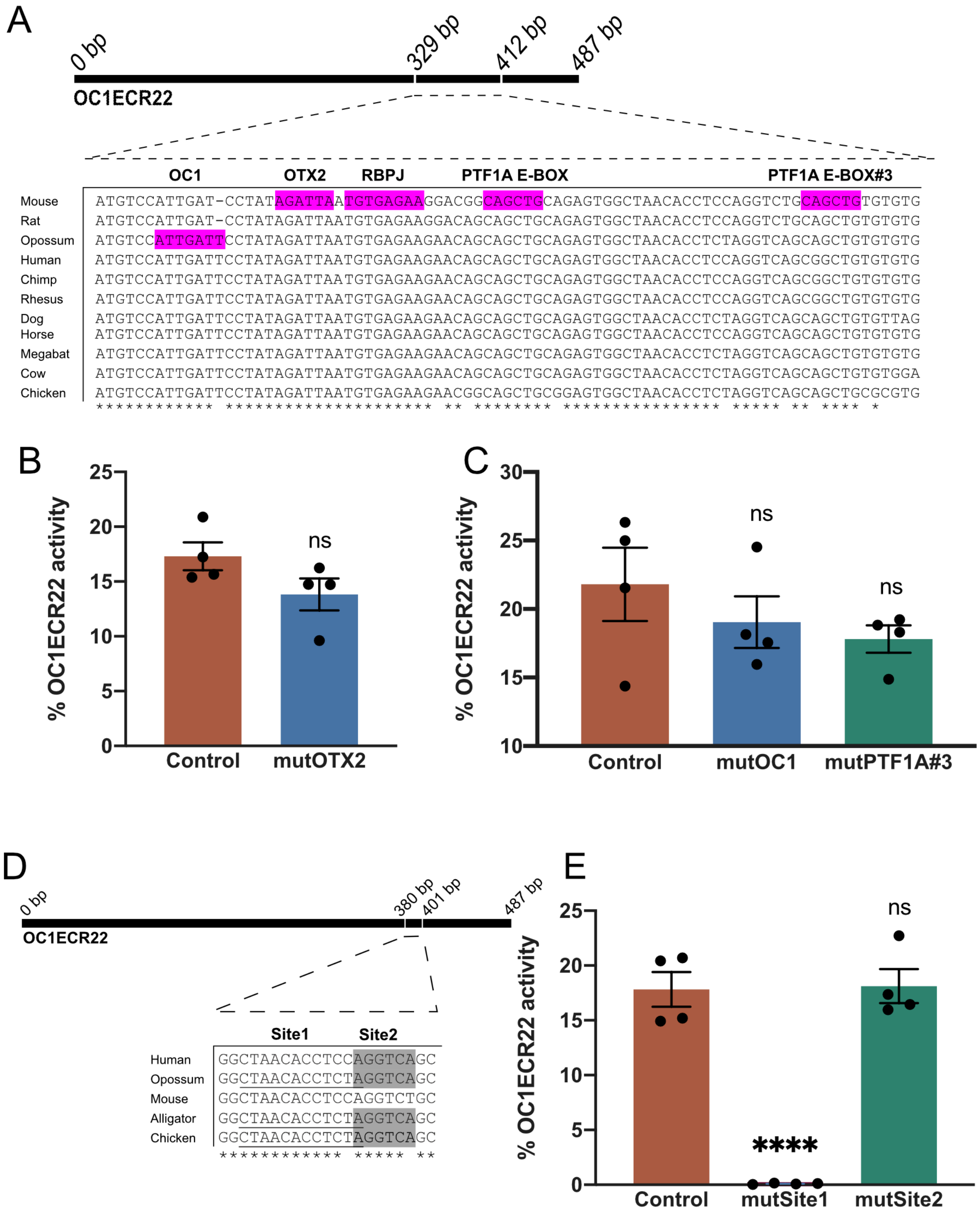

### Figure S6

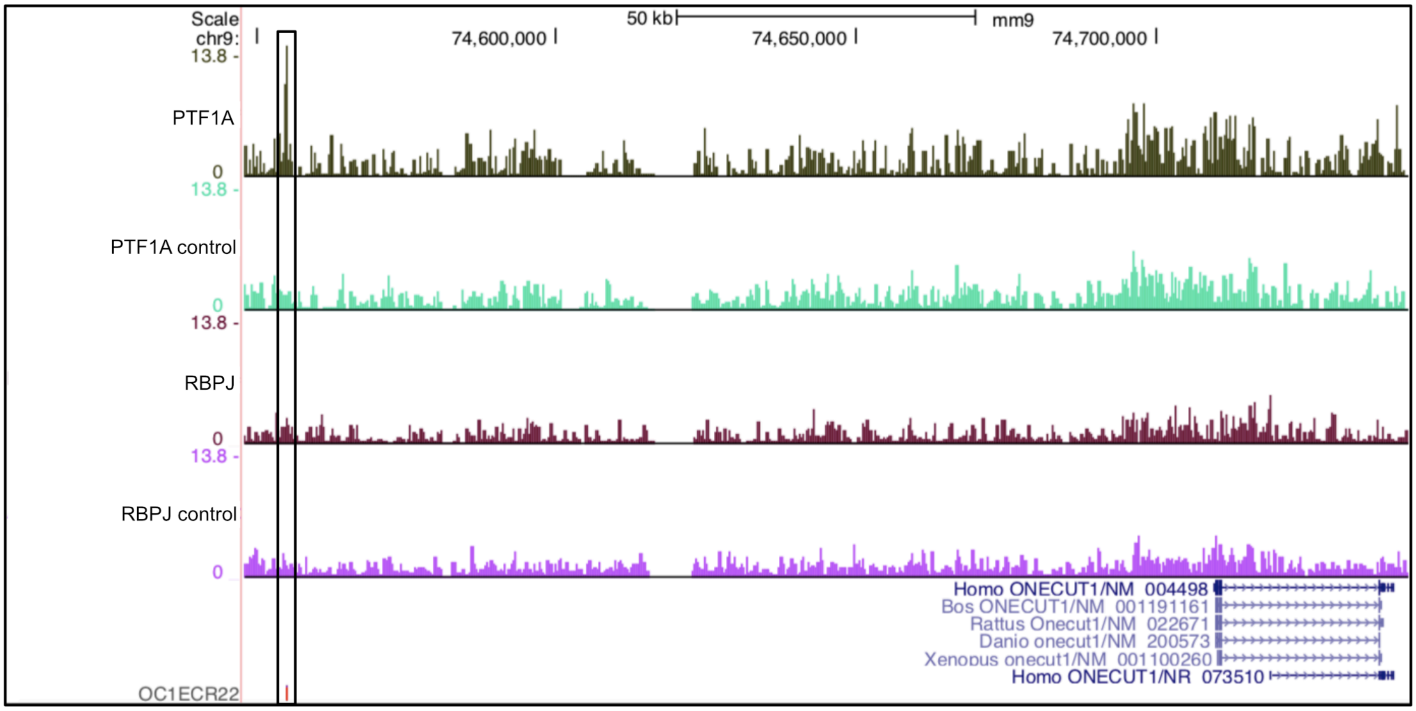

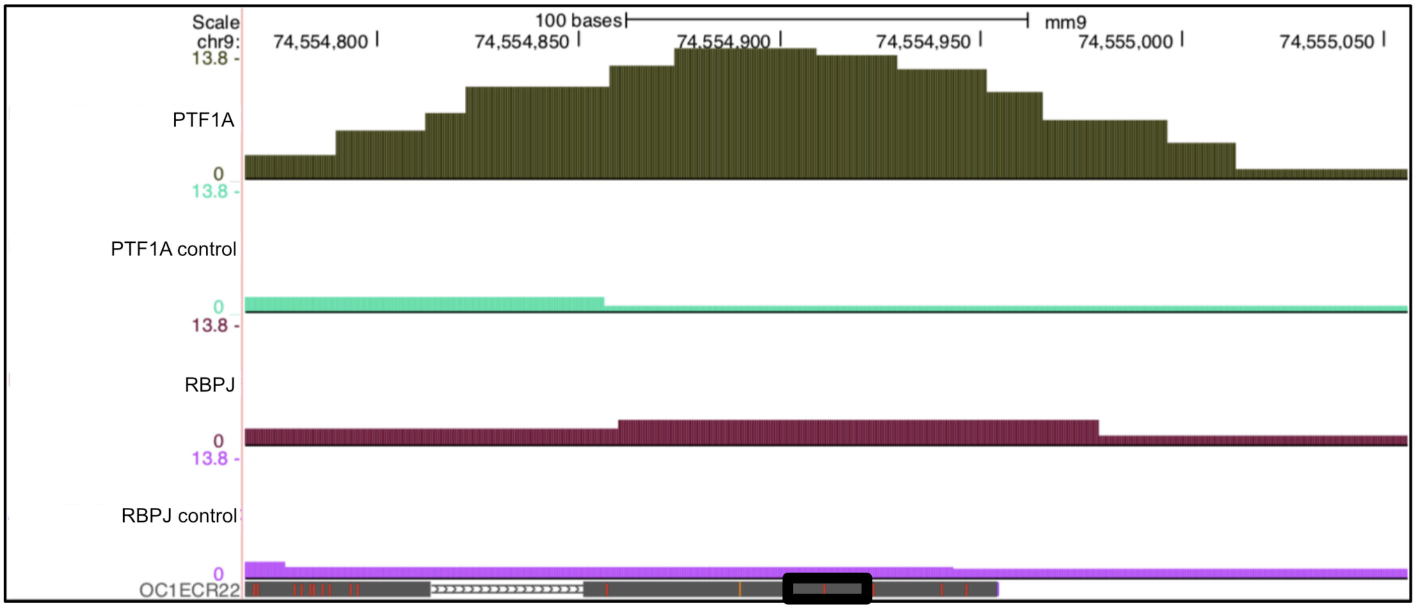

### Figure S7

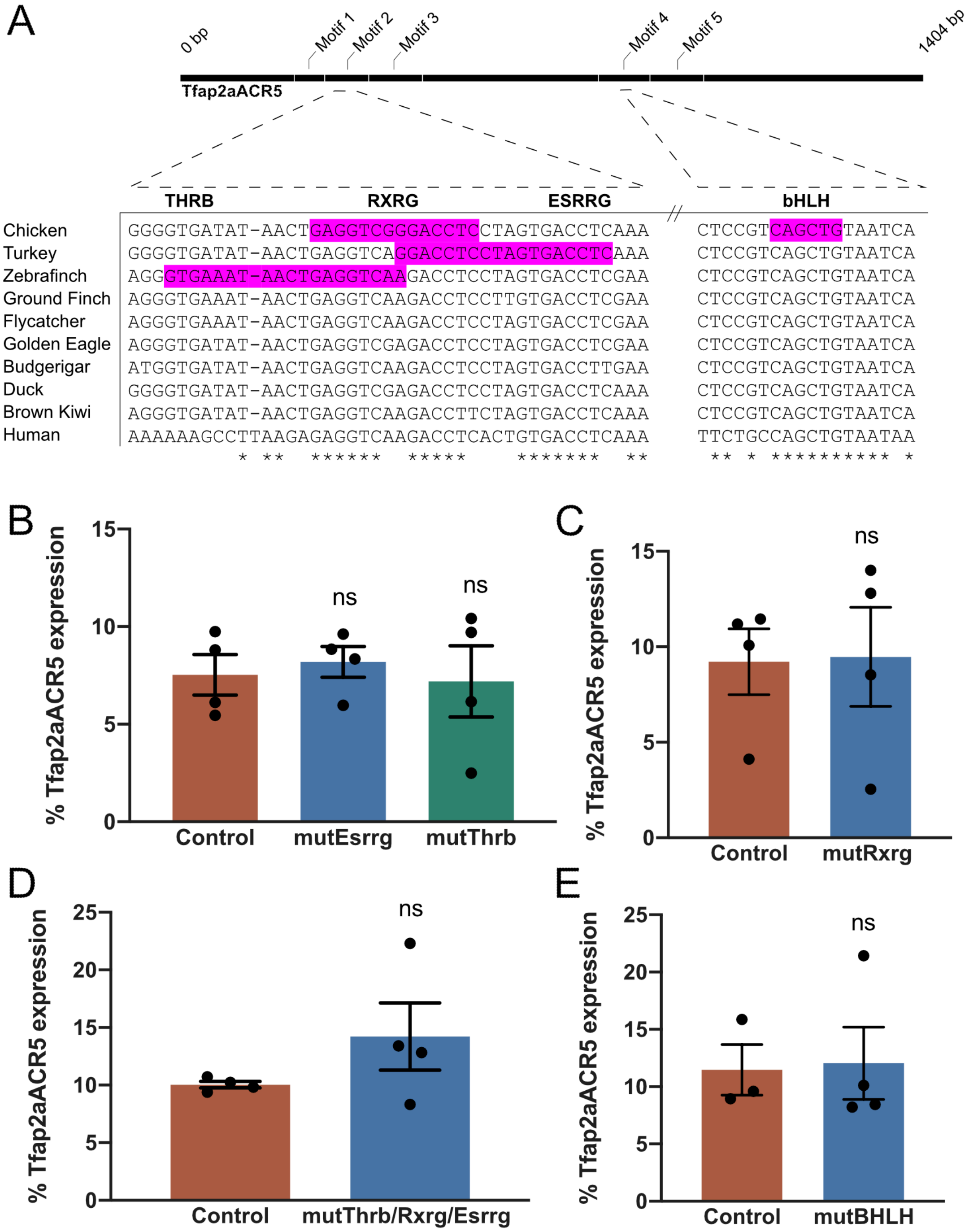
