## Supplementary material for "Early cis-regulatory events in the formation of retinal horizontal cells": Table 1

Active CRMs

| Enhancer Name | Source Genome | Genomic Coordinates | Alignment to Mouse Chain | Alignment to Human Chain |
|---|---|---|---|---|
| Ptf1aECR1 | mm10 | chr2:19,399,988-19,400,833 | n/a | yes |
| Ptf1aECR12[1] | galGal6 | chr2:17,407,144-17,407,747 | yes | yes |
| Ptf1aECR22 | galGal6 | chr2:17,378,344-17,378,735 | yes | yes |
| Ptf1aECR27[1] | mm10 | chr2:19,448,388-19,449,588 | n/a | yes |
| Ptf1aACR1 | galGal6 | chr2:17,444,278-17,444,694 | no | no |
| Ptf1aACR2 | galGal6 | chr2:17,393,167-17,393,644 | no | no |
| Ptf1aACR3 | galGal6 | chr2:17,367,462-17,368,314 | no | no |
| Ptf1aACR4[1+2] | galGal6 | chr2:17,414,463-17,415,436 | yes | yes |
| Ptf1aACR5 | galGal6 | chr2:17,417,914-17,418,243 | yes | yes |
| Ptf1aACR7 | galGal6 | chr2:17,506,117-17,506,589 | no | no |
| Lhx1ACR1 | galGal6 | chr19:8,532,231-8,533,204 | yes | yes |
| Lhx1ACR2 | galGal6 | chr19:8,465,565-8,466,343 | yes | yes |
| Lhx1ACR4 | galGal6 | chr19:8,515,541-8,516,138 | yes | yes |
| Tfap2aACR2 | galGal6 | chr2:63,308,785-63,309,676 | yes | yes |
| Tfap2aACR4 | galGal6 | chr2:63,440,976-63,441,857 | yes | yes |
| Tfap2aACR5 | galGal6 | chr2:63,411,881-63,413,275 | yes | yes |
| OC1ECR22[3] | galGal6 | chr10:9,057,472-9,057,958 | yes | yes |
| OC1ACR4[3] | galGal6 | chr10:8,978,219-8,979,121 | no | no |
| OC1ACR8[3] | galGal6 | chr10:8,857,680-8,858,311 | yes | yes |

Inactive CRMs

| Enhancer Name | Source Genome | Genomic Coordinates* | Alignment to Mouse Chain | Alignment to Human Chain |
|---|---|---|---|---|
| Ptf1aECR2[4] | galGal6 | chr2:17,421,529-17,422,001 | yes | yes |
| Ptf1aECR4 | mm10 | chr2:19,440,262-19,440,685 | n/a | yes |
| Ptf1aECR7[1] | galGal6 | chr2:17,412,272-17,412,743 | yes | yes |
| Ptf1aECR9[1] | galGal6 | chr2:17,409,243-17,410,082 | yes | yes |
| Ptf1aECR10[1] | galGal6 | chr2:17,408,534-17,409,159 | yes | yes |
| Ptf1aECR11[1] | mm10 | chr2:19,457,521-19,458,178 | n/a | yes |
| Ptf1aECR13[1] | galGal6 | chr2:17,406,726-17,407,164 | yes | yes |
| Ptf1aECR14[1] | mm10 | chr2:19,460,017-19,460,917 | n/a | yes |
| Ptf1aECR15 | galGal6 | chr2:17,405,574-17,406,029 | yes | yes |

| | | | | |
|---|---|---|---|---|
| Ptf1aECR16 | mm10 | chr2:19,477,634-19,478,022 | n/a | yes |
| Ptf1aECR17 | mm10 | chr2:19,484,196-19,484,363 | n/a | yes |
| Ptf1aECR18 | galGal6 | chr2:17,385,878-17,386,360 | yes | yes |
| Ptf1aECR19 | galGal6 | chr2:17,385,726-17,386,077 | yes | yes |
| Ptf1aECR20 | galGal6 | chr2:17,381,466-17,381,888 | yes | yes |
| Ptf1aECR23 | mm10 | chr2:19,520,397-19,521,218 | n/a | yes |
| Ptf1aECR24 | mm10 | chr2:19,523,808-19,524,236 | n/a | yes |
| Ptf1aECR25 | mm10 | chr2:19,406,822-19,407,137 | n/a | yes |
| Ptf1aECR28 | mm10 | chr2:19,449,608-19,449,920 | n/a | yes |
| Lhx1ACR5 | galGal6 | chr19:8,377,465-8,377,686 | no | yes |
| Tfap2aACR1 | galGal6 | chr2:63,277,143-63,277,844 | no | no |
| Tfap2aACR3 | galGal6 | chr2:63,310,201-63,310,611 | no | no |

[1] This sequence aligns with the 3' PTF1A enhancer described in Meredith et al., 2009
[2] A portion of this sequence aligns with the chick version of Ptf1aECR27
[3] This sequence was described in Patoori et al., 2020
[4] This sequence aligns with the 5' PTF1A enhancer described in Meredith et al., 2009
*Sequences for some inactive CRMs are centered within the listed genomic coordinates
